# Complex structural variant visualization with SVTopo

**DOI:** 10.1101/2025.04.16.649185

**Authors:** Jonathan R Belyeu, William J Rowell, Juniper A Lake, James Matthew Holt, Zev Kronenberg, Christopher T Saunders, Michael A Eberle

## Abstract

Structural variants are genomic variants that impact at least 50 nucleotides and can play major roles in diversity and human health. Many structural variants are complex multi-breakpoint rearrangements that are difficult to comprehend with existing visualization tools. We present SVTopo, a tool to visualize complex structural variants with supporting evidence from high-accuracy long reads, in easily understood figures. We include examples of eleven categories of complex structural variants from seven human genomes. SVTopo shows breakpoint evidence in ways that aid reasoning about the impact of large, multi-breakpoint events such as inversions, translocations, and combinations of simpler structural variants.

## Background

Structural variants (SVs) are responsible for more base pairs of genomic variation than any other class of variant and play major roles in diversity, rare disease, and complex disease risk (1–3). SV research has progressed in recent years from a focus entirely on very large variations in copy number (4,5) to much more inclusive studies (6–8) that report deletions, duplications, translocations, insertions, and inversions as small as 50 bp. SVs that do not precisely conform to any of these definitions have been reported as complex SVs (1,2,9–11). These generally consist of combinations of rearrangements of multiple classic SVs, such as a duplication followed immediately by a deletion or an inverted duplication. A common type is the unbalanced inversion, where an inversion occurs in tandem with deletion or duplication of flanking genomic material, found to represent one fifth of inversions larger than 2 kbp (12).

Technologies such as optical mapping, nanopore sequencing, and single-molecule real-time sequencing that generate multi-kilobase read lengths allow individual reads to span multiple breakpoints in many SVs. New software tools for SV calling (13–16) and genome phasing (17,18) use this additional information to increase SV recall and identify relationships between breakpoints in complex SVs. However, as large numbers of increasingly complex SVs are identified, they highlight challenges in the essential next step of understanding SV impact on important genes, regulatory elements, and other factors in genomic health.

Visualization is a critical step in variant analysis for most variant types and several tools exist for variant visualization(19–21), allowing researchers to evaluate variant call evidence and review likely impacts on nearby genomic features. The distinct challenges in comprehending SVs identified from long reads have resulted in development of new tools for SV visual curation, differentiating real variants from false positives by visual inspection of supporting evidence (22,23). Tools for improved comprehension and reasoning about SVs have also been created, often with a focus on extremely large-scale variation such as is seen in tumor contexts (24,25). Despite the development of these tools there remains a gap for visualizing complex SVs, likely leading to missed insights of these variants on human health.

SVTopo was created to simplify and enhance localized complex SV visual inspection. The goal of SVTopo is to help humans comprehend complex SV effects on genome structure and impact on critical gene regions. SVTopo presents the essential read-alignment components of complex SV evidence in the context of individual rearranged genomic blocks, creating clear and high-resolution graphical representations of extremely complex rearrangements. The images aid in human reasoning about genomic impacts of complex SVs, assisting users in grasping potential results on human genomic health, gene structure, and regulatory elements. SVTopo improves researcher understanding of complex structural components of genomic topography.

SVTopo enhances analysis for many types of challenging variants. Multi-breakpoint SVs, inter- or intra-chromosomal translocations, inversions (with or without flanking insertions/deletions), and non-tandem inversions or duplications are all difficult to visualize with existing genome browsers, and thus to understand conceptually. SVTopo can be simply applied to genomic data and is readily added to genome analysis pipelines for small or large sequencing cohorts, greatly improving the accessibility of complex SV evaluation.

## Results

SVTopo uses chimeric alignments from phased high-accuracy sequencing to construct networks of connected genomic break-end locations. These networks annotate blocks of genomic material that are deleted, duplicated, inverted, relocated, or otherwise rearranged. SVTopo creates visual representations of the blocks, including their order, alignment support, and orientation in the sample genome. This provides a simple comparison between the sample and reference structures. For example, in **Figure 1** an SV with a structure difficult to understand via IGV or Ribbon visualization is intuitively represented by SVTopo. SVTopo can optionally annotate images with gene locations and other genomic information important to the user, such as locations of transposons or repeats. Image design carefully balances between simplicity and inclusion of essential details for complex SV interpretation, to enhance human understanding of the SV image.

**Figure 1.**
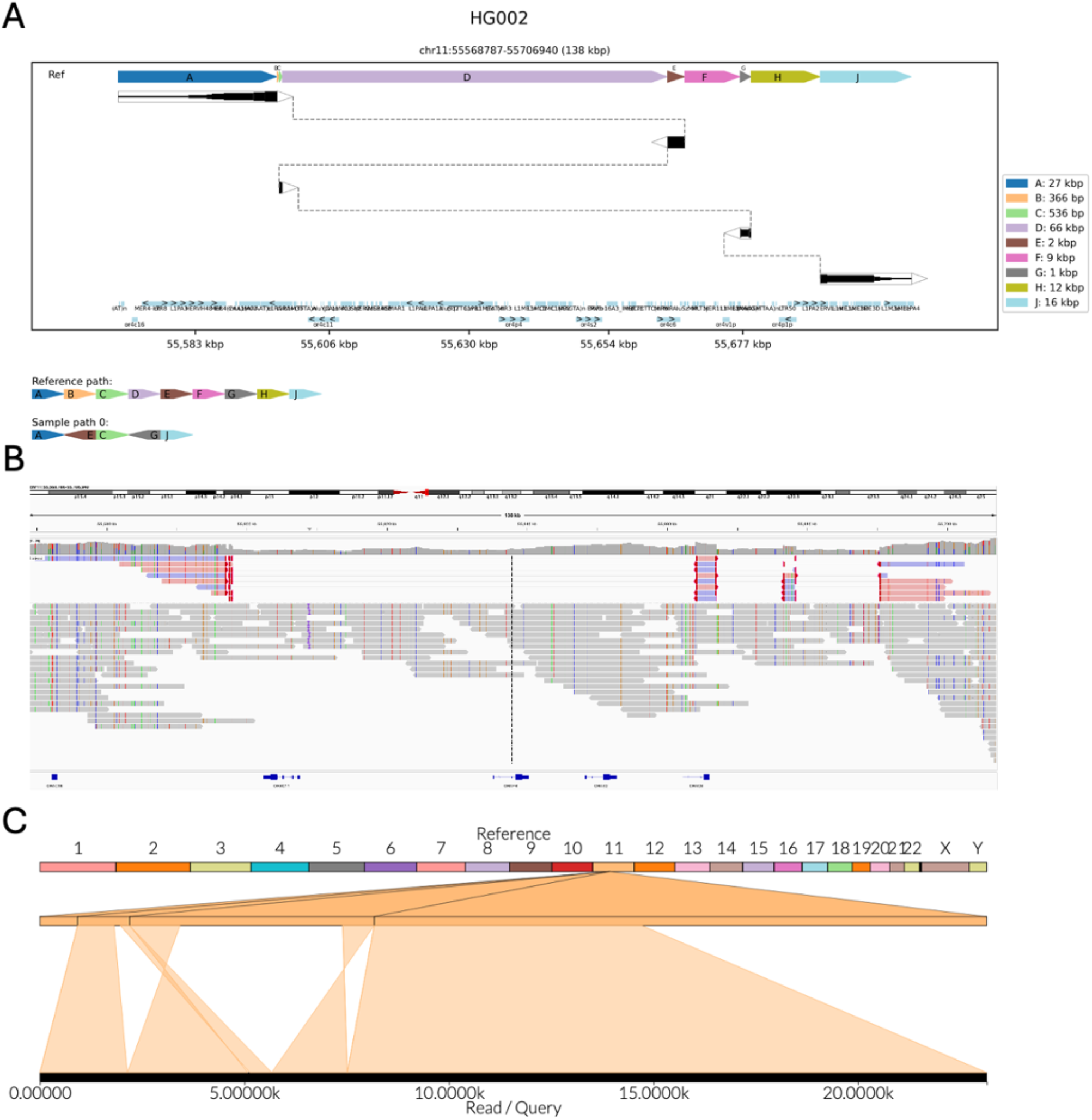
Visualization of a complex SV. Visual representations of an extremely complex rearrangement, consisting of deletions of four component blocks and inversions of two. **A**. An SVTopo plot clearly shows the order and orientation of inverted segments. Dark blocks represent alignment segments, made thicker where more alignment support occurs. Dotted lines connect blocks that are present in the sample. Blocks are ordered top-to-bottom as they appear in the sample genome and left-to-right as they appear in the reference genome. The bottom track below the main SV window contains a chain-plot of blocks in reference order, then sample order and orientation. Blocks B, D, F, and H in the Sample Path are eliminated, while C, E, and G are reordered. **B**. IGV representation of the region. **C**. Ribbon representation of the region.

### Multi-sample complex SV characterization

SVTopo was used to characterize complex SVs in a set of seven unrelated HiFi genomes (**see Methods**), generating 446 images. This includes 125 images with incompletely resolved structure; these indicate challenges in alignment and can in many cases be useful for identification of false-positive complex SV calls (see **Supplementary Figures 1-5**). 37 images represented simple pairs of deletion or duplication variants associated by phasing. The remaining 284 images represented high-confidence complex SVs. Many of these were shared by multiple samples, with a total of 142 unique complex SV loci (26). While well-defined categories do not exist for most complex SVs, these variants were manually assigned to eleven definitions ranging from relatively simple to extremely complex (see **Figure 2** and **Supplementary Table 1**). Five singleton SVs were individually unique and were grouped together as Unique-singleton variants, with the other ten categories containing structurally similar SVs.

**Figure 2.**
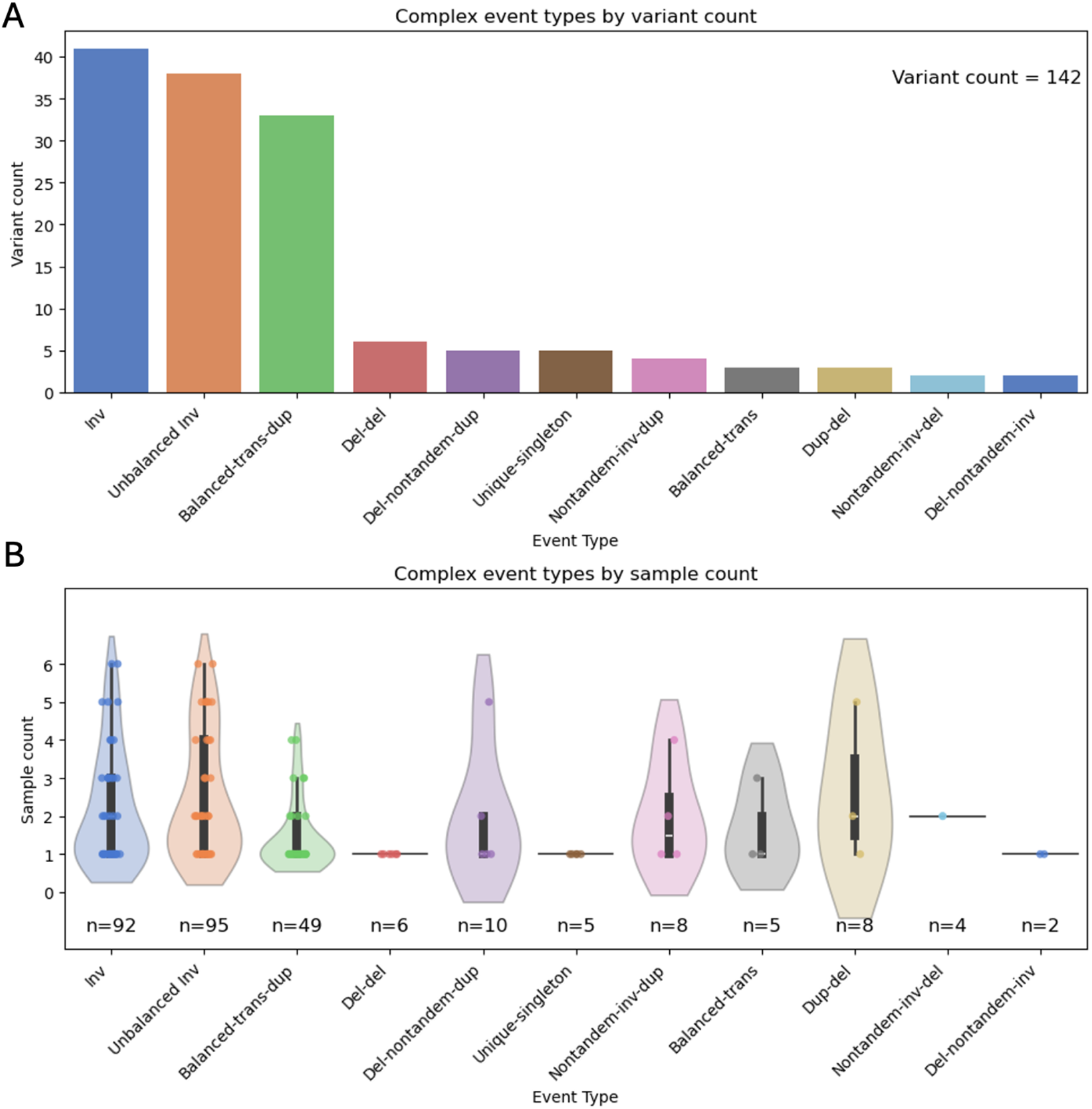
Types and distributions of complex SVs in seven unrelated genomes. 142 unique complex SVs were manually assigned to eleven categories corresponding to variations on simple SV types, with five complex SVs grouped as Unique-singleton. **A**. Counts of unique SVs assigned to each category. **B**. Distributions of sample counts for variants belonging to each SV category.

Many of these complex SVs render very simply with SVTopo but are extremely challenging to visualize with existing tools. This is demonstrated by examples of non-tandem duplications and balanced translocations with flanking deletions, as shown in **Figure 3**. See also **Supplementary Figures 6-12** for examples of additional complex SV types with comparison to IGV and Ribbon visualizations.

**Figure 3.**
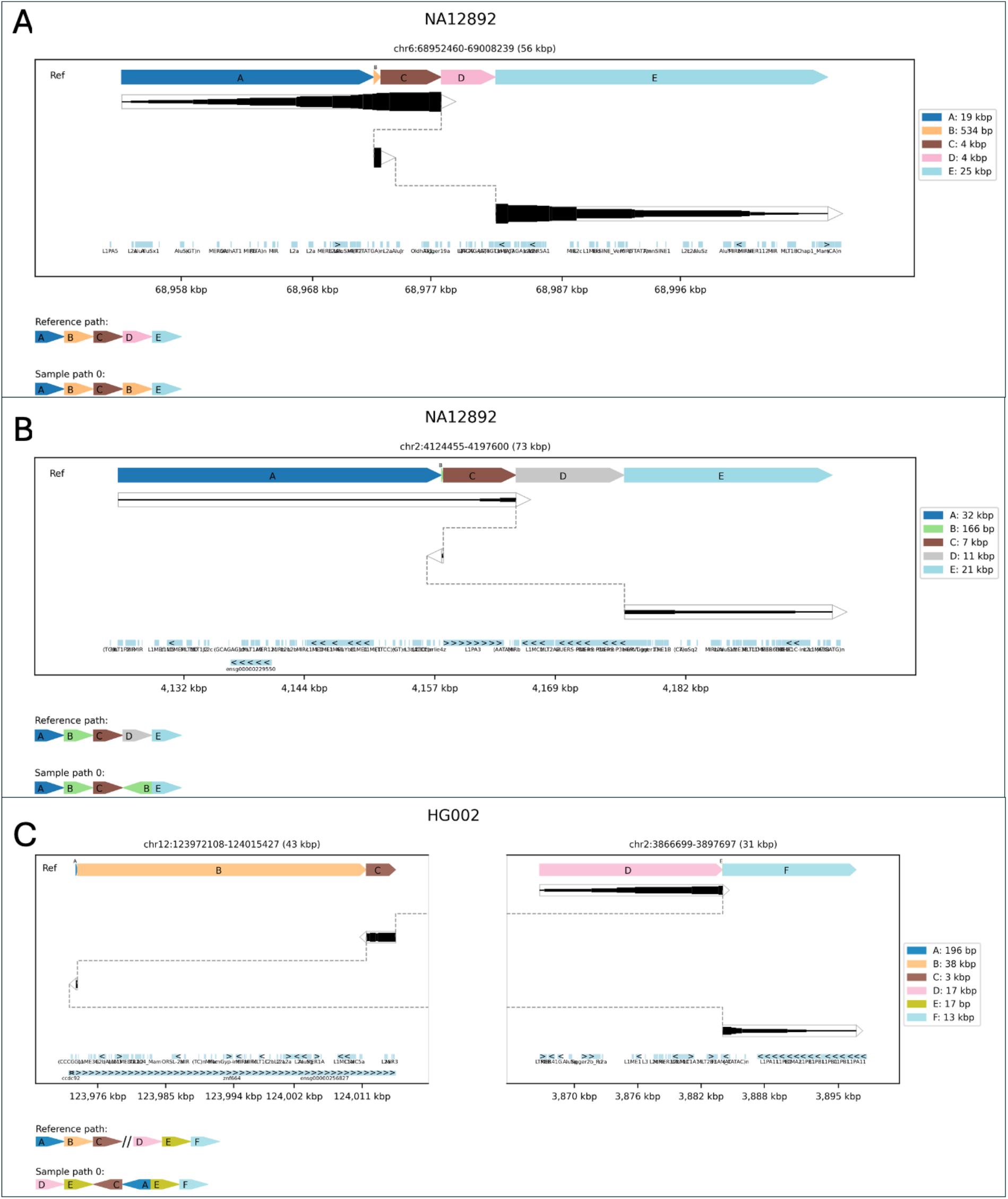
Examples of three complex SV categories. Plots of three complex SV types that are challenging to interpret without SVTopo representations. **A**. Dup-del: non-tandem duplication followed by a deletion. **B**. Nontandem-inv-del: An inverted non-tandem duplication followed by a deletion. **C**. Balanced-trans-dup: A balanced translocation consisting of two blocks from chr12 translocated to chr2 and the duplication of a short region on chr2.

The most common type of SV in this analysis were inversions, which were grouped into three categories: balanced inversions (Inv, n=44), with a genomic block reversed in orientation and directly reconnected; inversions with exactly one flanking deletion (Inv-del, n=17); and inversions with flanking deletions on both sides (Inv-double-del, n=25). Visualizations of these variants can be quite challenging using standard techniques, as visualizations not implemented for complex SVs struggle to show each of the component rearrangements distinctly (See **Figure 4**).

**Figure 4.**
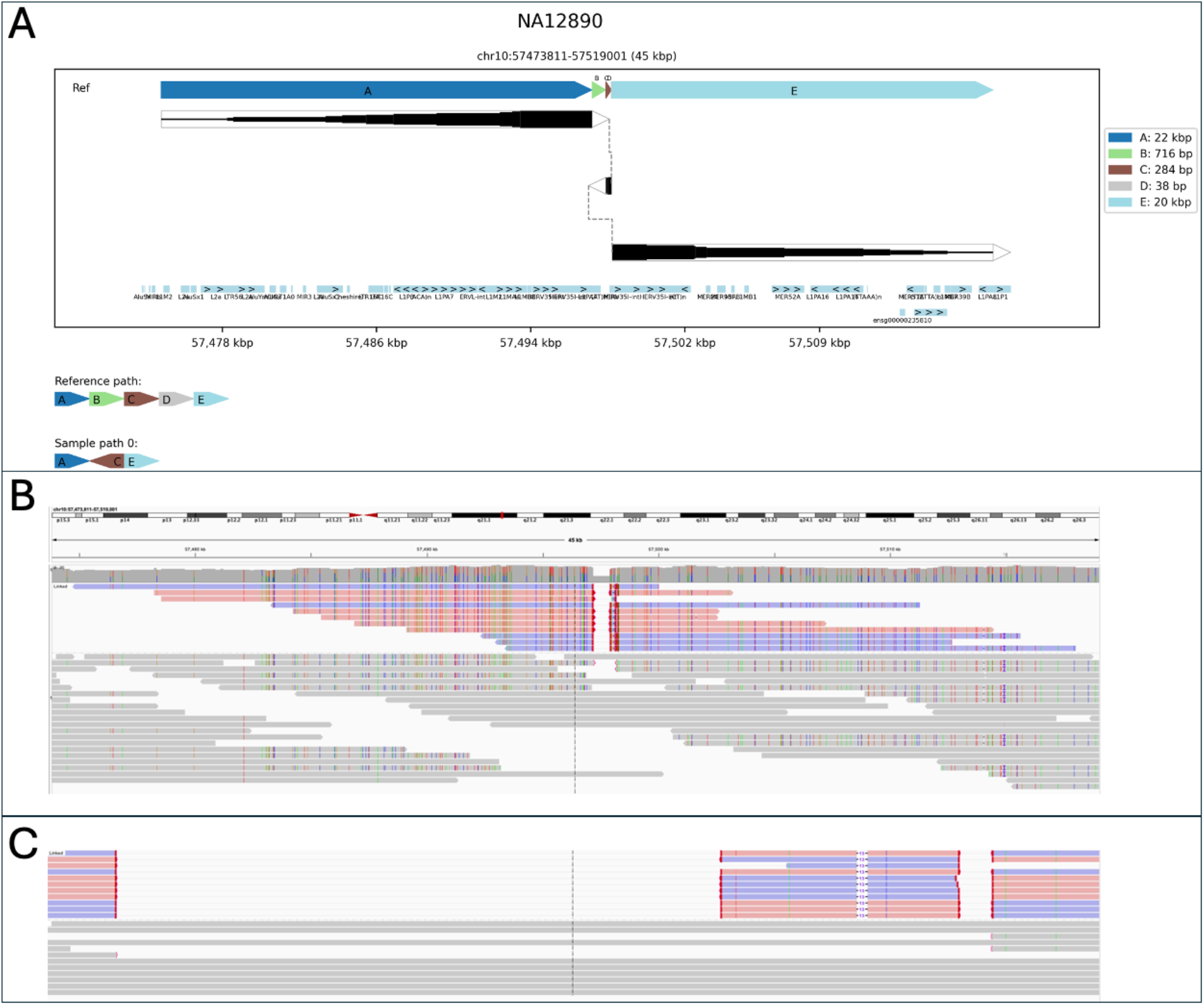
Example complex inversion with flanking deletions. A 284 bp inversion flanked by a 716 bp deletion and a 38 bp deletion (category Del-inv-del). **A**. SVTopo plot of region. **B**. IGV screen capture of the region with linked supplementary alignments. **C**. IGV image of the region, zoomed in to include only complex SV breakpoints.

A total of 44 unique balanced inversion variants were found, many occurring in multiple samples to appear a total of 83 times. 17 distinct inversions with a single flanking deletion were found, appearing a total of 42 times; and 25 with two flanking deletions, appearing 52 times. Many of the inversions were accompanied by large flanking deletions, often larger than the sizes of the inverted blocks (see **Figure 5** and **Supplementary Table 2)**.

**Figure 5:**
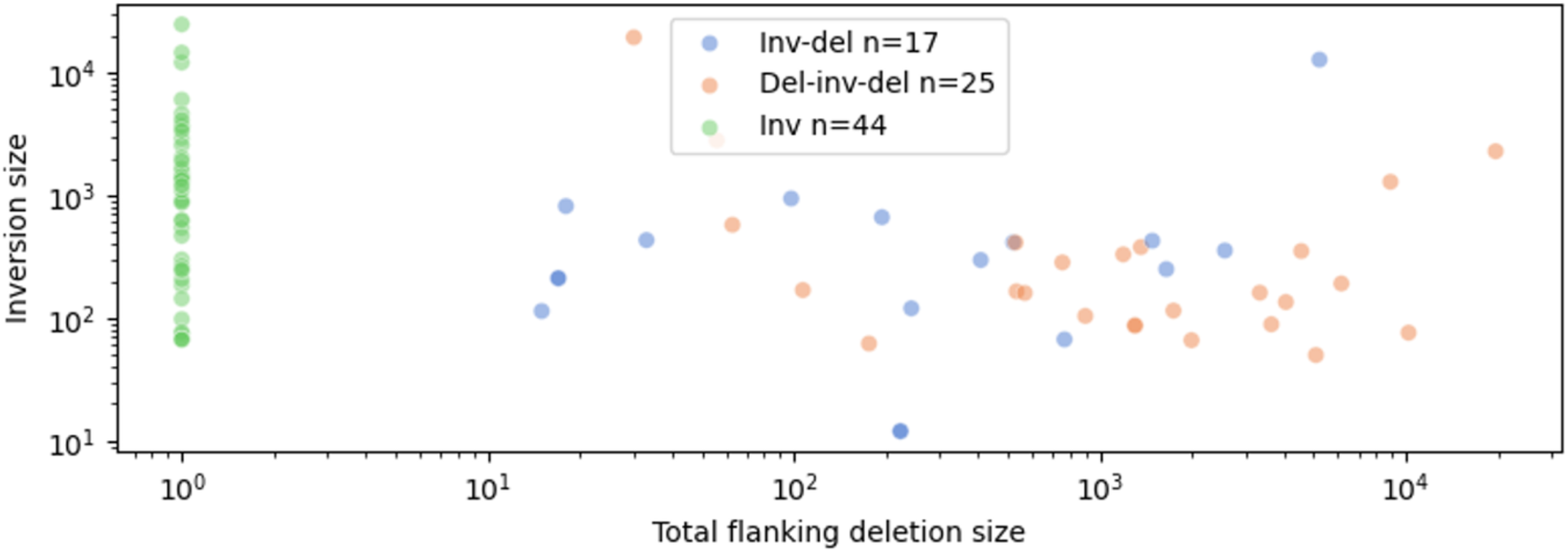
Balanced and unbalanced inversion sizes. Lengths of inverted blocks contrasted against flanking deletion sizes for inversions with 0-2 flanking deletions. Each axis is shown on log scale. Non-tandem inversions excluded.

### SVTopo table viewer

In addition to providing simple visualizations for complex SV review, SVTopo automatically creates a feature-rich serverless HTML-based viewing table to organize and present images (see **Figure 5**). This viewer is a self-contained web page, with built-in filtering tools to allow users to rapidly identify the complex SVs most relevant to a research priority (such as reviewing images relevant to a region or sample of interest). The viewer can be easily deployed locally for a single user or remotely if multi-user access is desired (example viewer publicly available (27)). The viewer main page consists of a variant region table, with a row for each genomic window represented. Clicking a row displays the image. Columns in the table show the chromosome, start, end, region size, associated variant IDs (if available), number of variant IDs represented in the image (if available), and sample ID. The table allows a user to filter by clicking a variant ID, number of variants field, or sample ID. A filtering options window also allows filtering on region size ranges and target chromosomes.

## Discussion

SVTopo provides a new toolkit for visual analysis of complex SVs, with an easy-to-understand plot layout and simple deployment interface. Analysis of seven unrelated genomes revealed 142 unique complex SVs in eleven categories. These demonstrate the utility of this tool due to the variety and frequency of complex SVs that call for specialized visualization approaches. SVTopo will be especially useful for 1) understanding the complexity of SV calls in whole genome analysis, with total per-genome complex SV counts that remain tractable for human review, and 2) reviewing genome structure in specific regions with known complex SVs.

These workflows will dramatically improve the facility of complex variant review versus existing tools and improve the ability of researchers to rapidly gain an understanding of hard-to-characterize rearrangements, along with improved understanding of impact on important genes. SVTopo can be used as a standalone tool or in tandem with existing tools such as IGV, Ribbon, etc., enabling multiple levels of visual review for complex SVs.

The unique design of SVTopo images improves complex SV interpretation, conveying information with individual blocks of genomic material shown with both sample and reference order. Explicit block orientations and connections between blocks inform the user of how genomic regions are rearranged, essential for understanding potential impacts on genetic disease from novel adjacencies and lost or duplicated material. Examples of easily interpretable SVTopo representations of complex rearrangements in **Supplementary Figures 6-12** show these advantages. In balanced translocations, for example (**Supplementary Figure 6**), existing tools may mask small flanking duplications or make it difficult to identify and view donor sites. Similarly, the vast array of possible non-tandem inversions and duplications are extremely challenging to understand with existing tools (see **Supplementary Figures 8-12**), as showing such rearrangements with a focus on either reference or sample ordering of blocks can mask the relationships between the two. Variations of an inversion between two deletions (**Figure 4**) can be difficult to differentiate from a pair of deletions around a non-inverted region (**Supplementary Figure 7**). These examples demonstrate the necessity of using dedicated software to understand localized complex SVs, to fully interpret consequences on genomic structure and health.

These results show that it is not uncommon for a genome to contain multiple distinct types of complex rearrangements. As shown in **Figure 2**, these seven genomes included 142 unique complex SV loci with eleven distinct rearrangement types. The sample frequency of some complex SVs, appearing in multiple unrelated genomes even within this small cohort, indicates likely high population frequencies and increases their potential for impacts on evolutionary diversity, and possibly on complex disease. Inversions, often considered a simple SV type, are shown here to occur nearly as often with flanking deletions as without (44 unique balanced inversions vs 42 with flanking deletions). This finding also demonstrates how better variant visualization can improve our understanding of the effects of inversion on genomes and of the mutational mechanisms that drive structural rearrangement.

The SVTopo table viewer, designed to enhance genome analysis for small and large projects or teams, is a significant addition to the SVTopo toolkit. The ease of deployment and filtering options provides necessary assistance to researchers for identification of complex SVs relevant to important genomic questions and improve the utility of SVTopo for sharing conclusions among teams or collaborators.

## Conclusions

SVTopo makes extremely complex rearrangements simple to understand and interpret. It increases the utility of high accuracy long reads to more fully demonstrate the complexity of the genome, recovering insights into highly complex genomic rearrangement that otherwise escape notice or human comprehension. SVTopo performs complex reconstructions necessary to reveal the structure of extremely complex rearrangements in simple and human-accessible ways.

## Methods

### Defining the structure of complex variation with SVTopo

SVTopo uses highly accurate long-read alignments to identify breaks in a sample genome, connects these breaks into networks of regions that are rearranged relative to the reference genome, annotates them with supporting alignment counts, and filters regions with abnormal coverage or mapping quality prior to reporting complex SV events (see **Figure 7**). Resulting complex SV information is written to file on a per-sample basis and used to inform automated generation of descriptive complex SV plots. A central HTML webpage is generated for one or more samples processed together. For algorithm details, see **Supplementary Note 1**.

**Figure 6.**
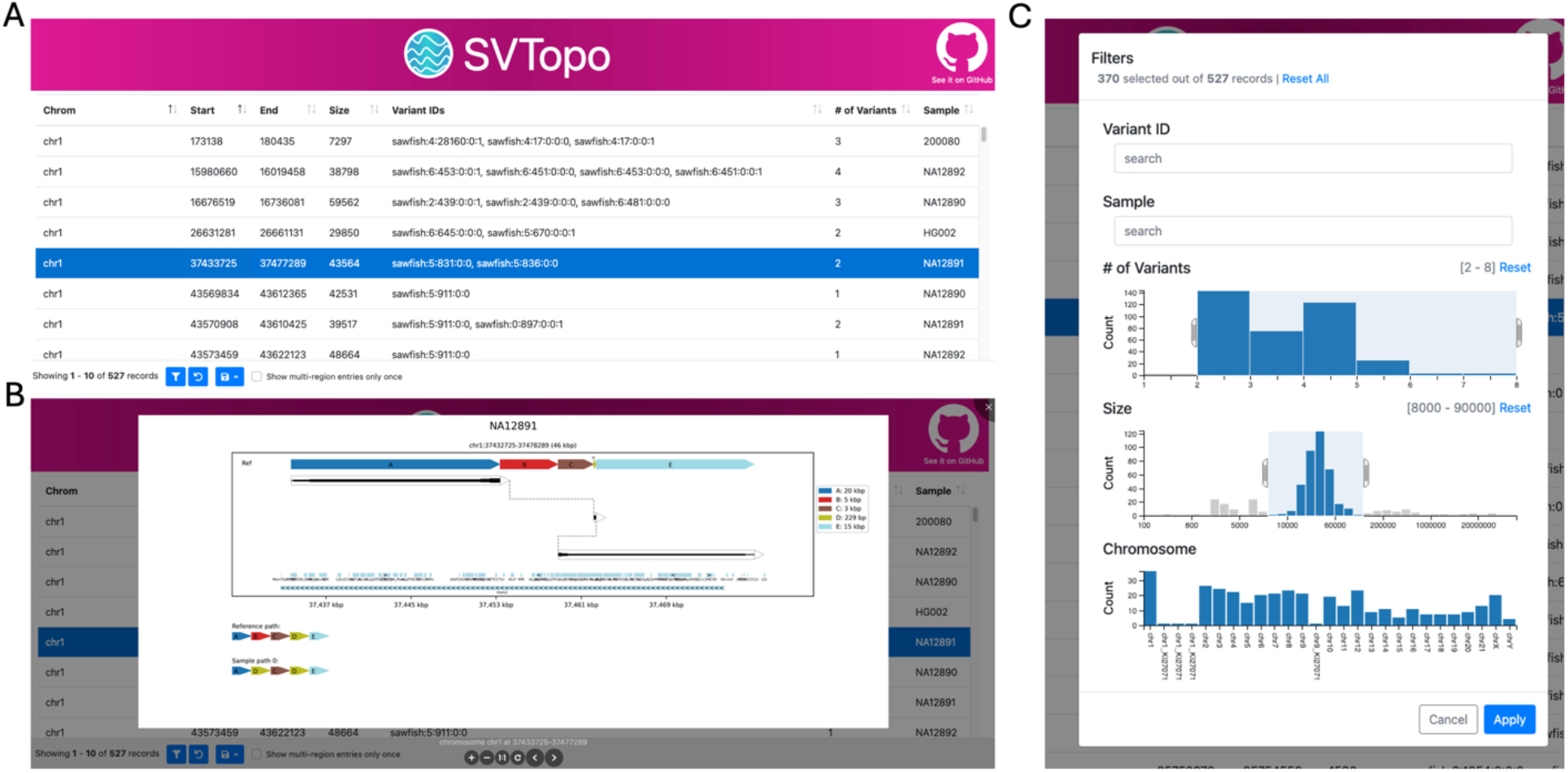
SVTopo table viewer. **A**. A screen capture of the table viewer, showing included table columns. **B**. A screen capture of a complex SV image displayed by clicking a table row. Image shows a deletion and non-tandem duplication. **C**. Viewer filtering window, with filtering fields for variant ID, sample ID, number of variants, region size, and chromosome. Filters are selected for 2-8 variants per image, a size range of 8,000-90,000 bp, and all chromosomes.

**Figure 7.**
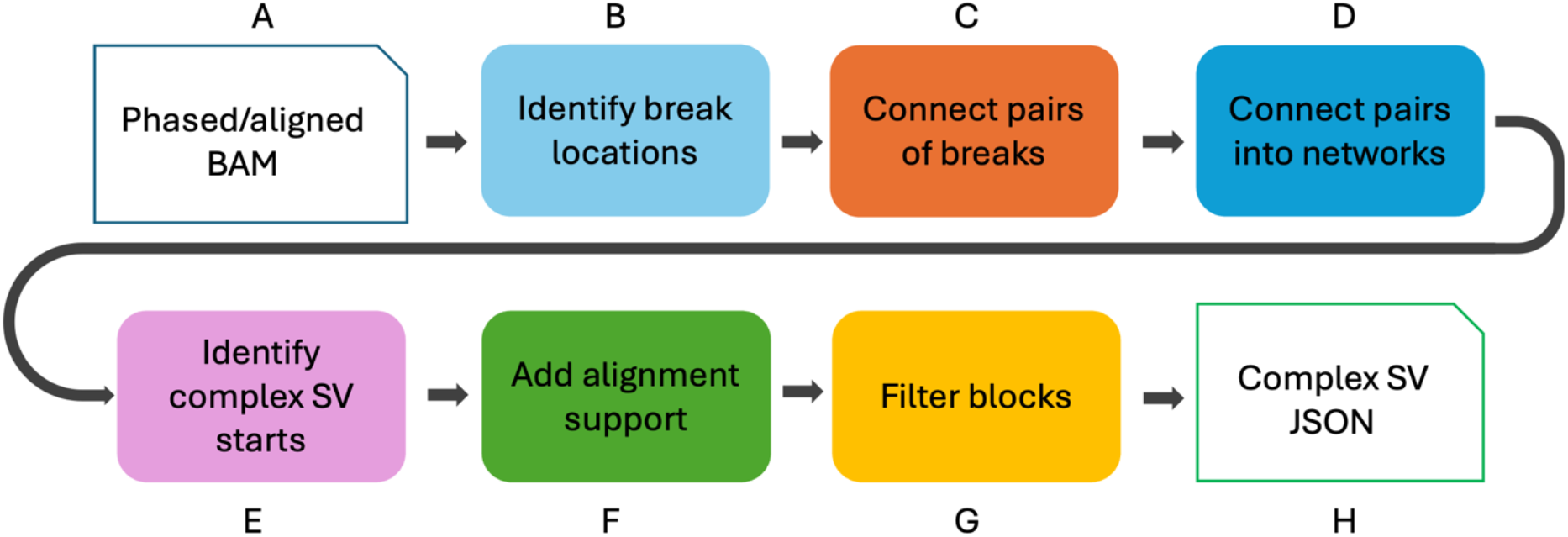
Identification of complex SVs with SVTopo. **A**. SVTopo uses phased and aligned BAM files of long reads as input, optionally accepting variant call information from Sawfish as well. **B**. Break locations are identified using read clipping locations, assisted by SV call coordinates if available. **C**. Read alignments and phase blocks are used to connect break locations together, forming pairs of connected breaks. **D**. Pairs of breaks are further connected into networks of connected break locations, forming full complex SV representations. **E**. Start breaks are identified for each network of connected break locations, allowed the identification of the correct event order in the sample. **F**. The number of supporting alignments for each pair of connected breaks in complete SV events is counted and added. **G**. Filters eliminate regions with abnormal coverage or highly repetitive sequence. **H**. Results are stored in JSON format, with additional annotations in BED format. These are used to inform automatic plot generation with SVTopoVz.

### Analysis of complex SVs in seven unrelated genomes

SVTopo was used to characterize complex SVs in a set of seven HiFi genomes: Genome in a Bottle Ashkenazi Trio Son HG002 (28) and six publicly released genomes from the three-generational Platinum Pedigree cohort (29). These six genomes (NA12889, NA12890, NA12891, NA12892, 20080, and 20100) were the unrelated parental genomes in that cohort, thus representing all included ancestral haplotypes in the Platinum Pedigree. All genomes were aligned to the GRCh38 human reference genome and phased with WhatsHap v1.4. SV calling was performed with Sawfish v0.12.7. Sample coverage depth was approximately 30x for HG002 and 200080, 45x for 200100 and NA12890, and 60x for NA12889 and NA12892. SVTopo identified complex SVs and generated images for all samples on a single processor in 8 hours, 39 minutes, about 1.25 hours per genome. The maximum RAM utilization was 4.75 GB.

