## Supplementary Figures for "Complex structural variant visualization with SVTopo"

### Slide 1
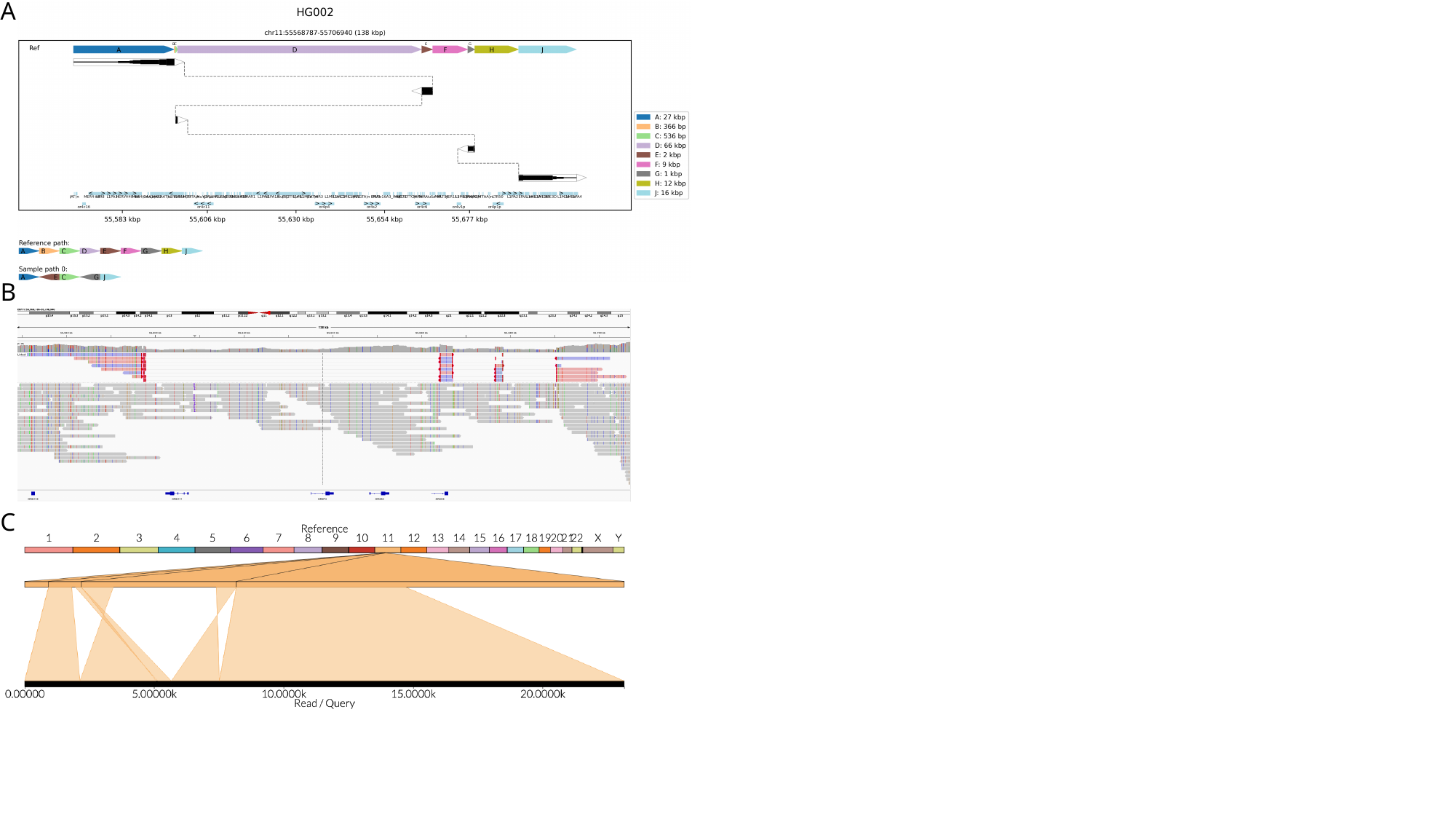

A
B
C

### Slide 2
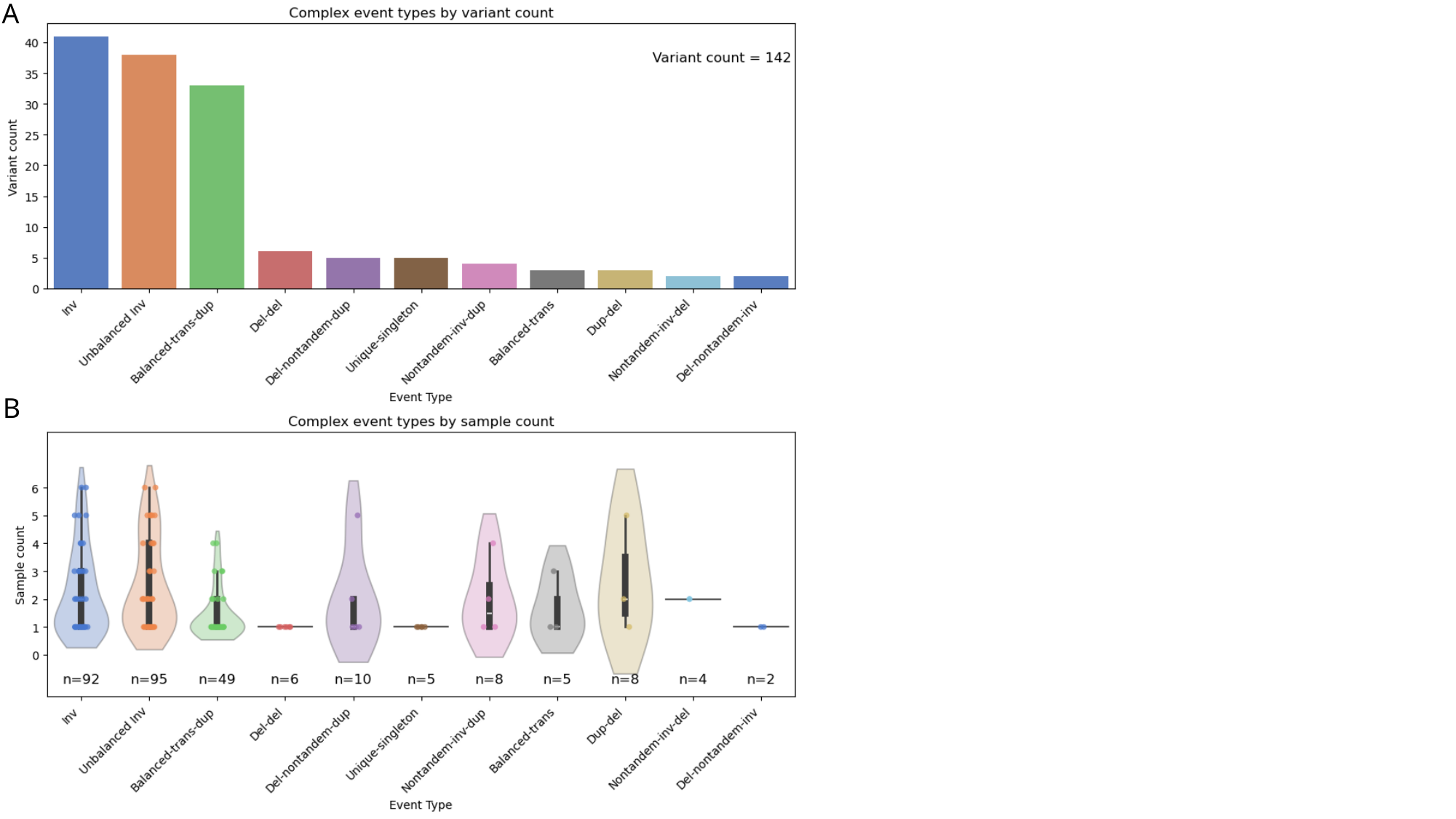

A
B

### Slide 3
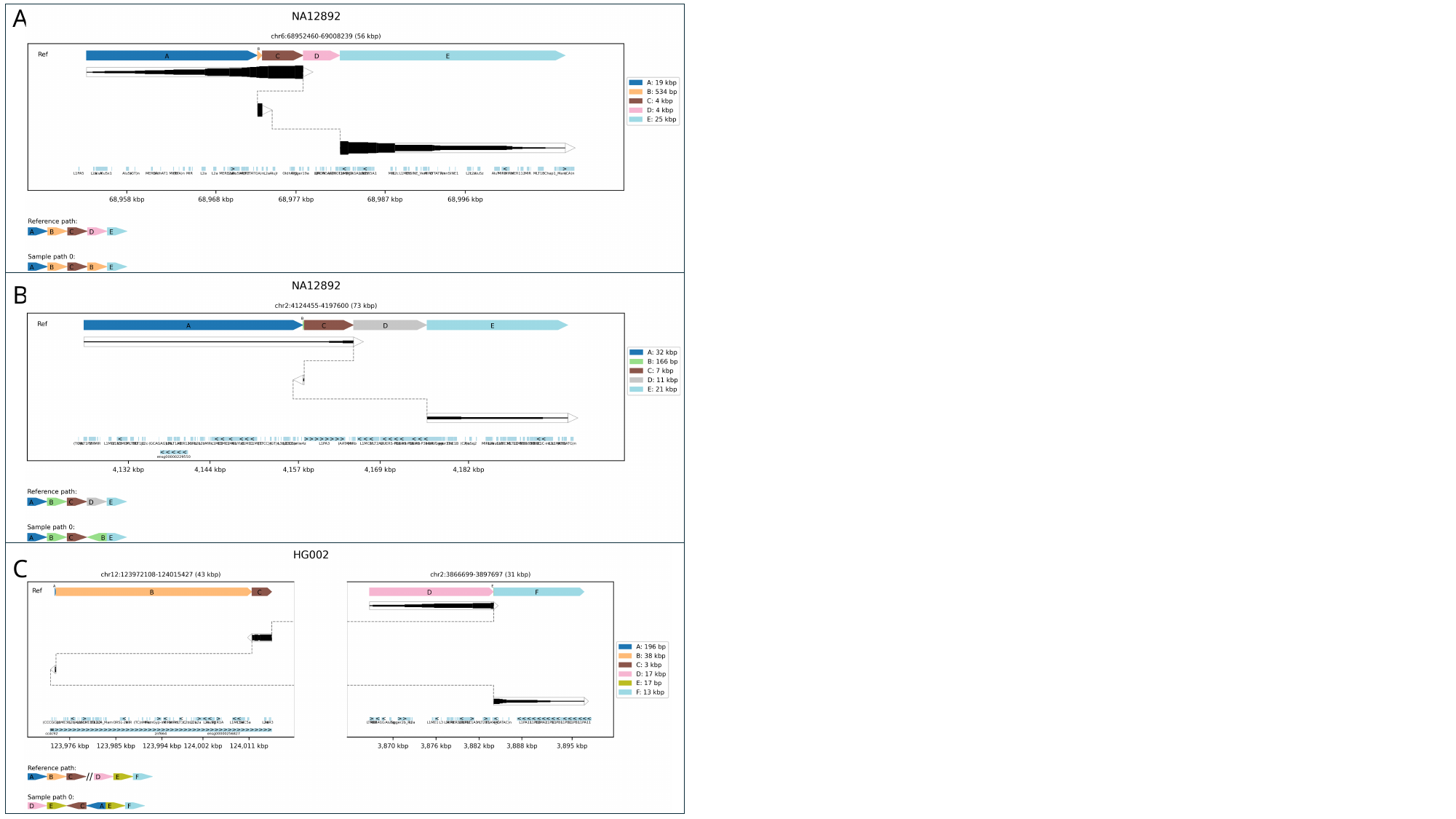

A
B
C

### Slide 4
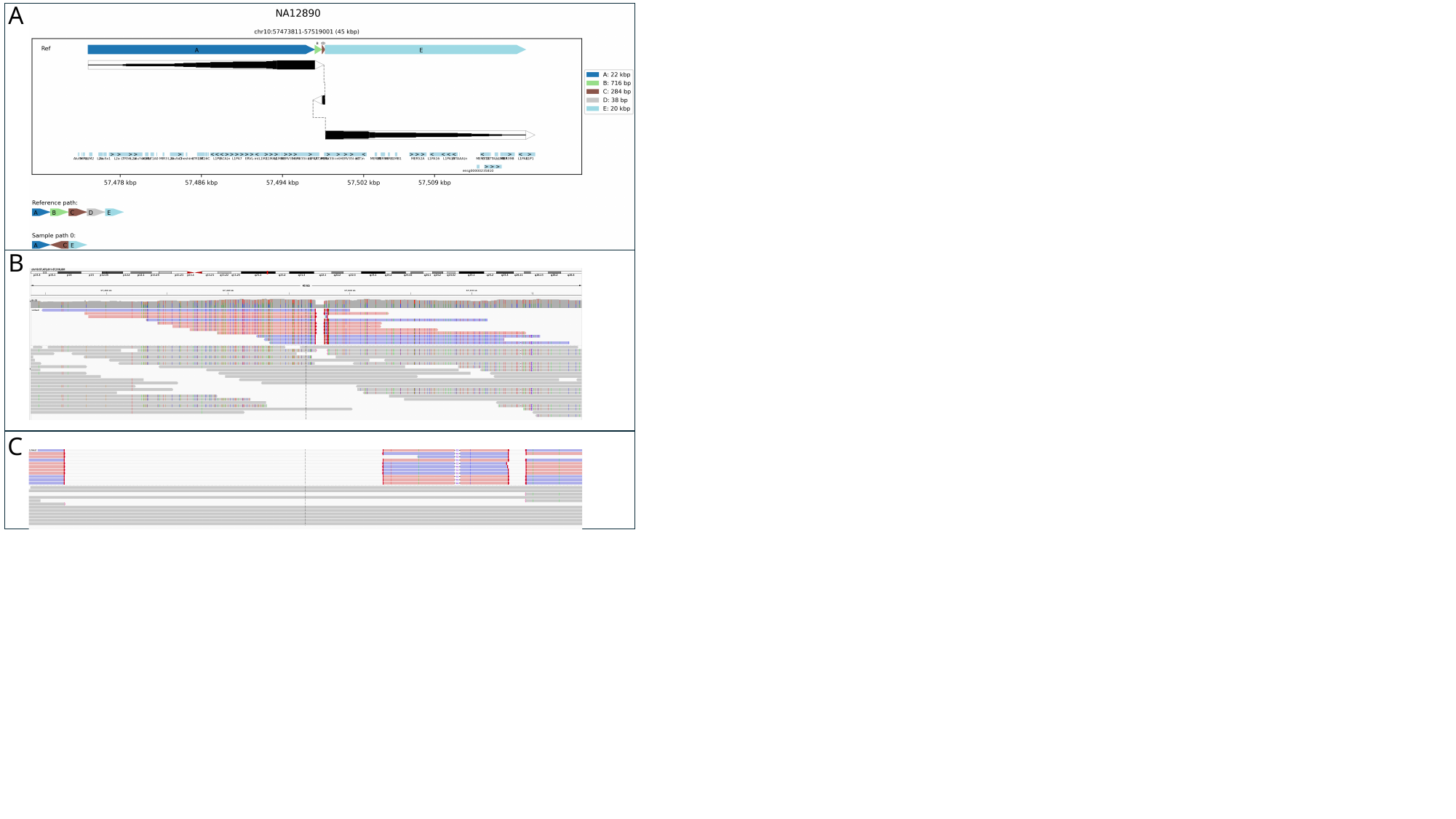

A
B
C

### Slide 5
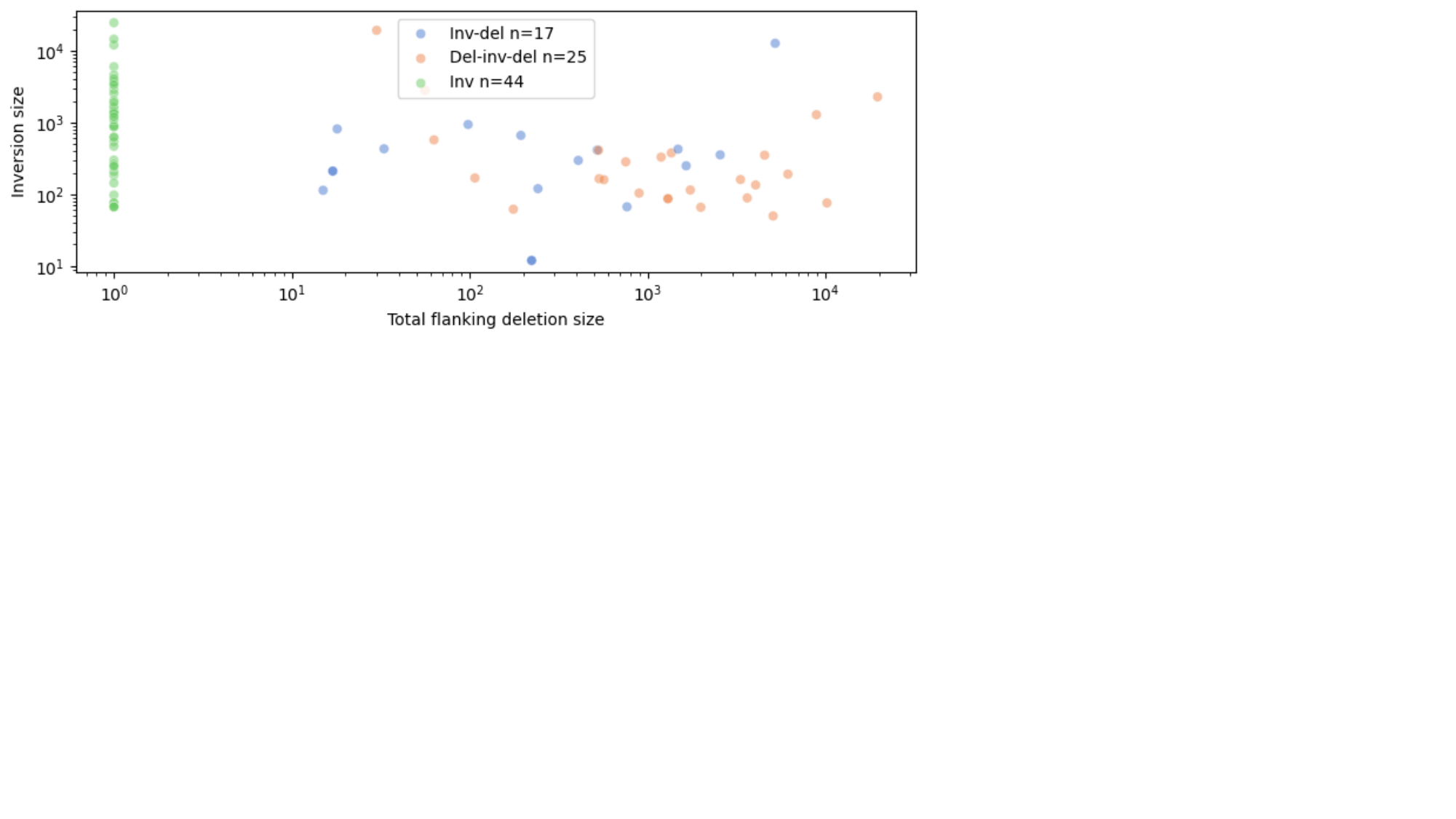

### Slide 6
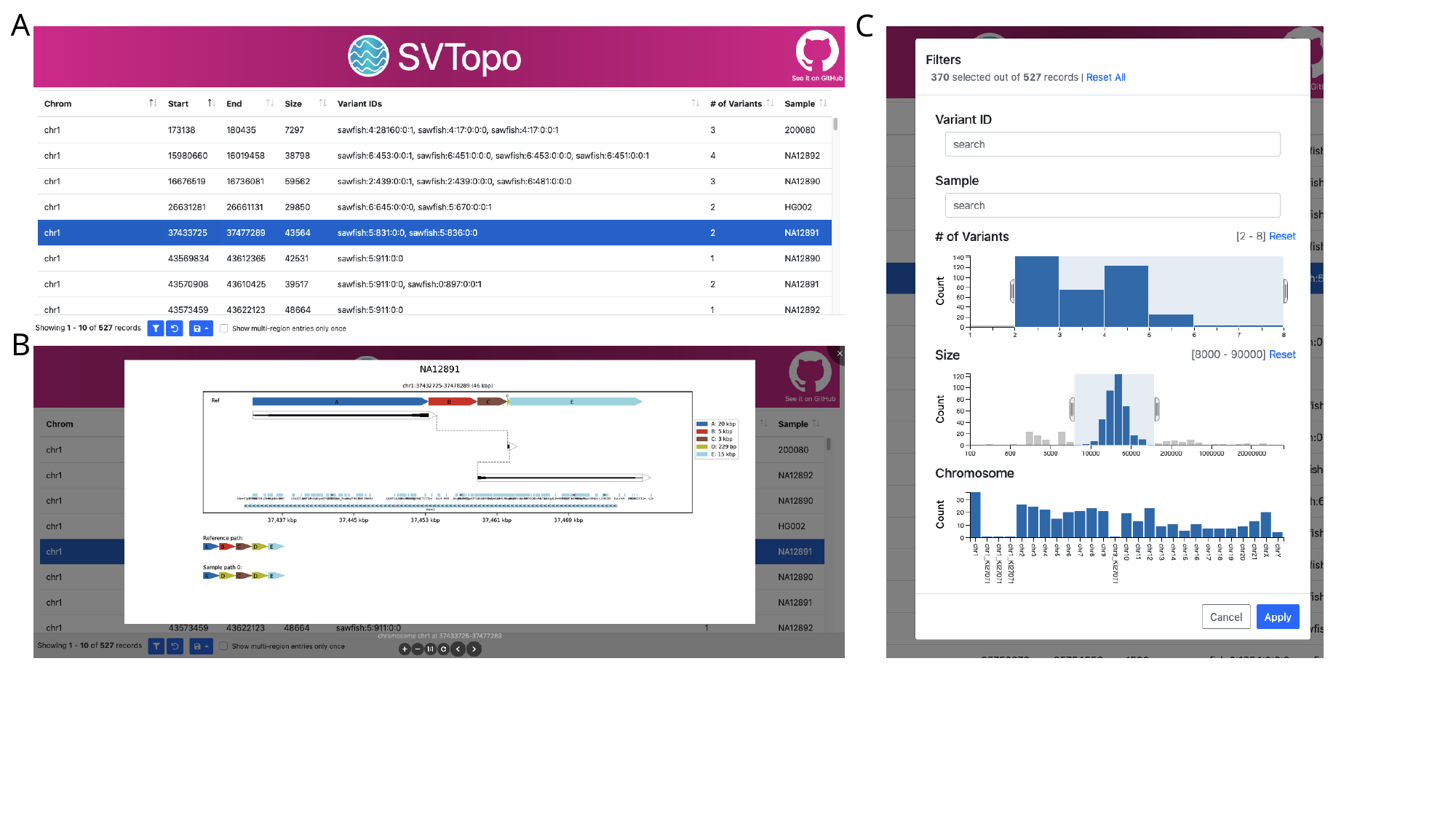

A
C
B

### Slide 7
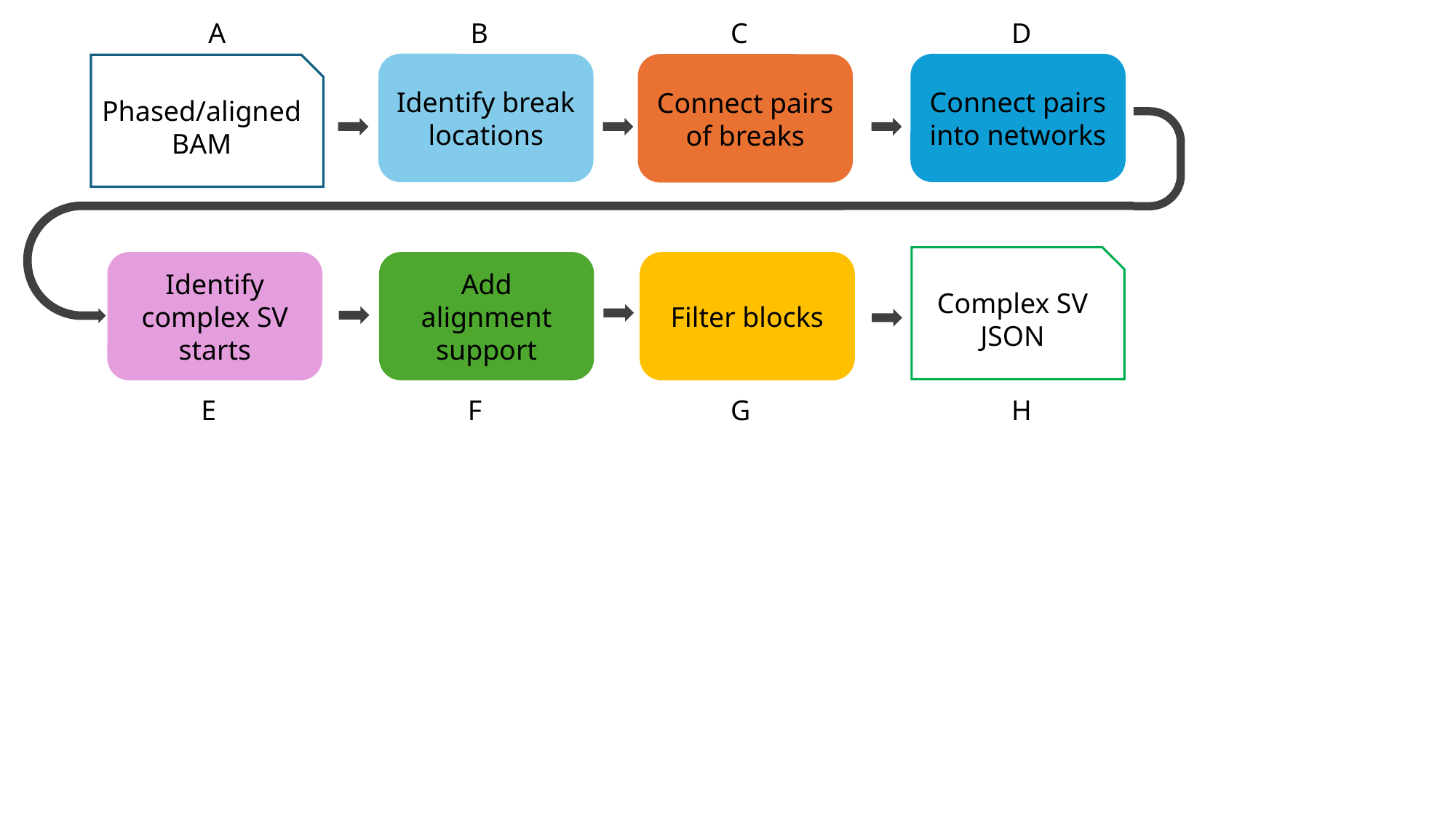

A
B
C
D
Phased/aligned BAM
Identify break locations
Connect pairs into networks
Connect pairs of breaks
Complex SV JSON
Identify complex SV starts
Add alignment support
Filter blocks
E
F
G
H

### Slide 8
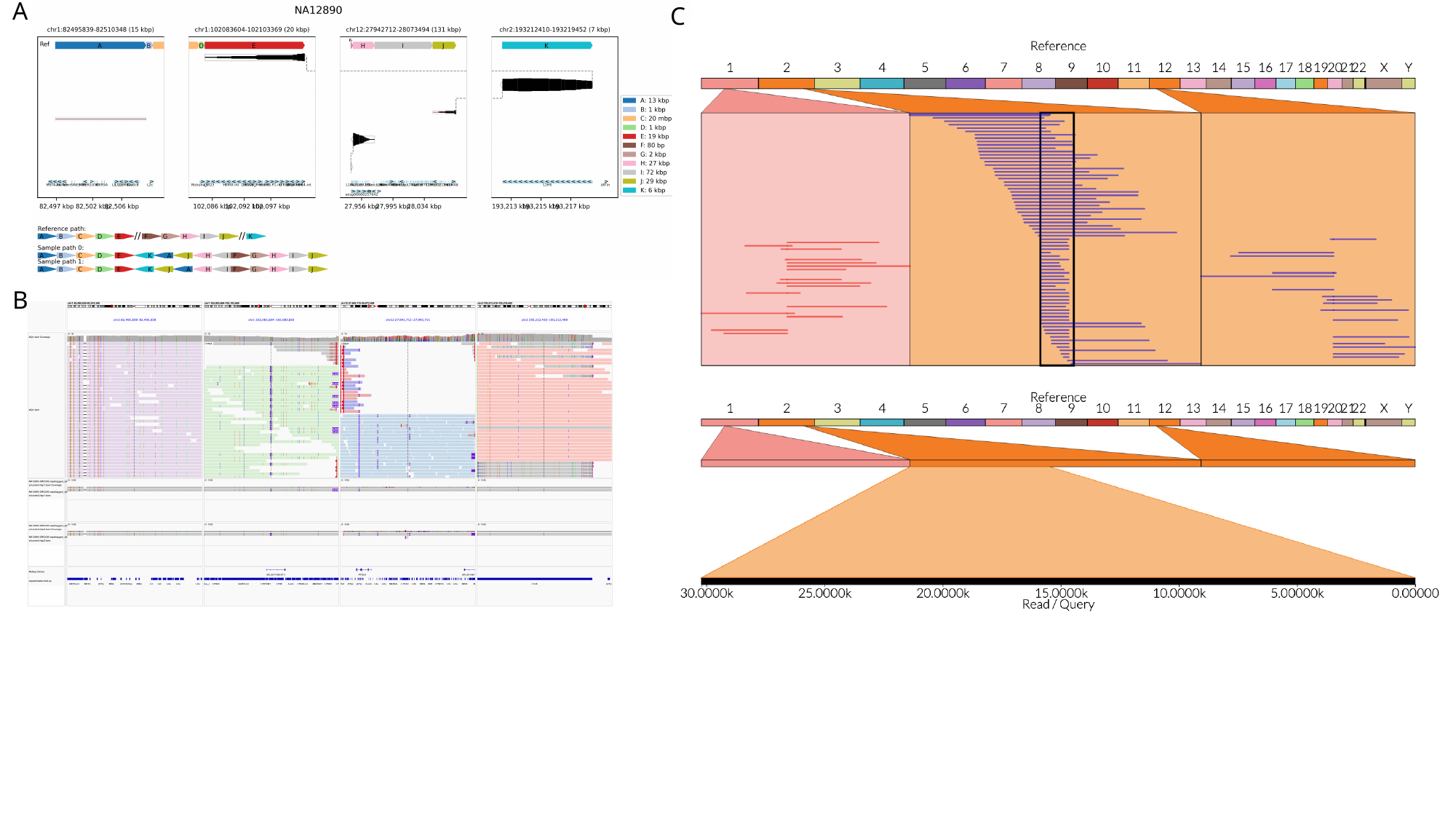

A
C
B

### Slide 9
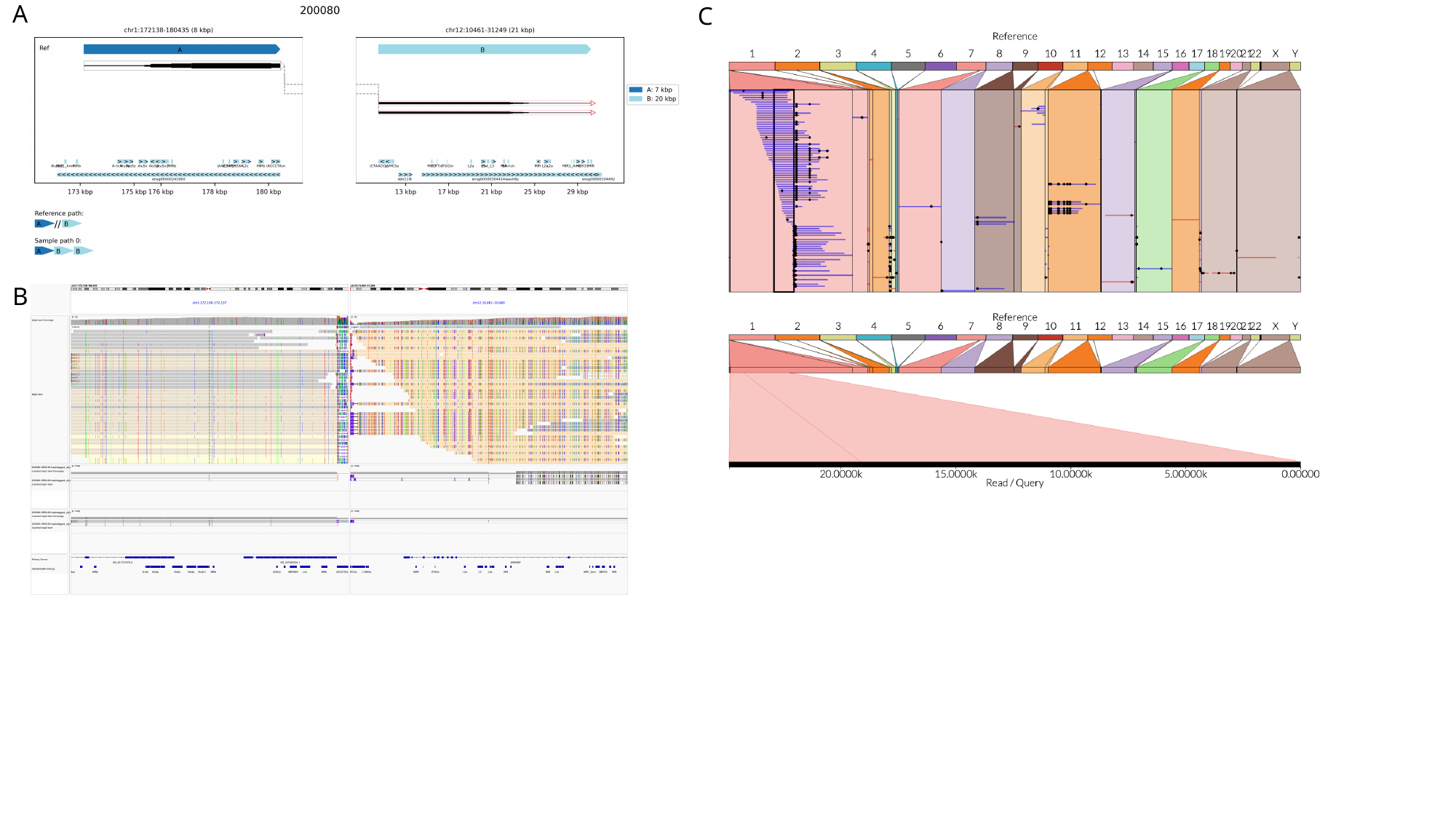

A
C
B

### Slide 10
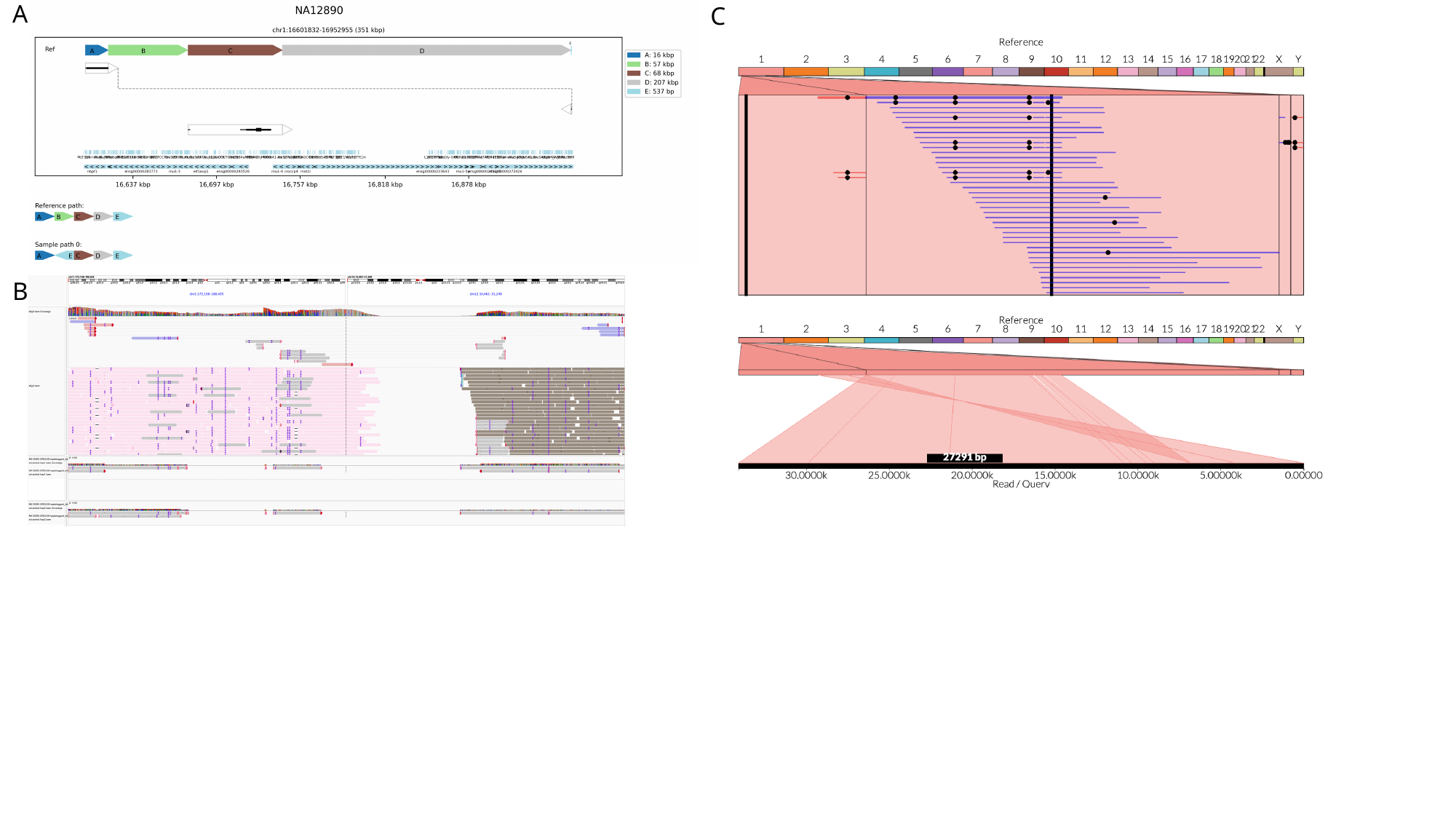

A
C
B

### Slide 11
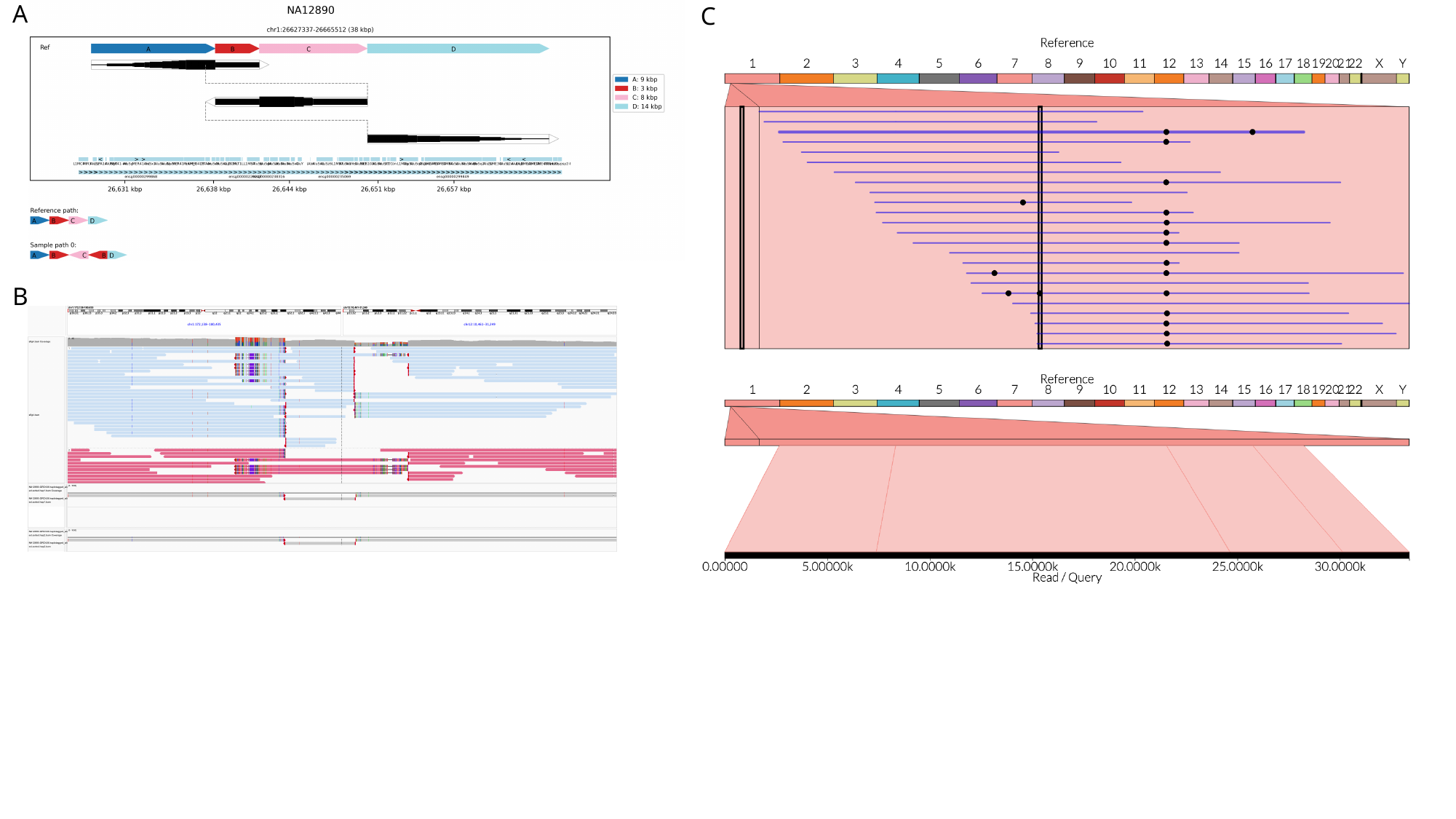

A
C
B

### Slide 12
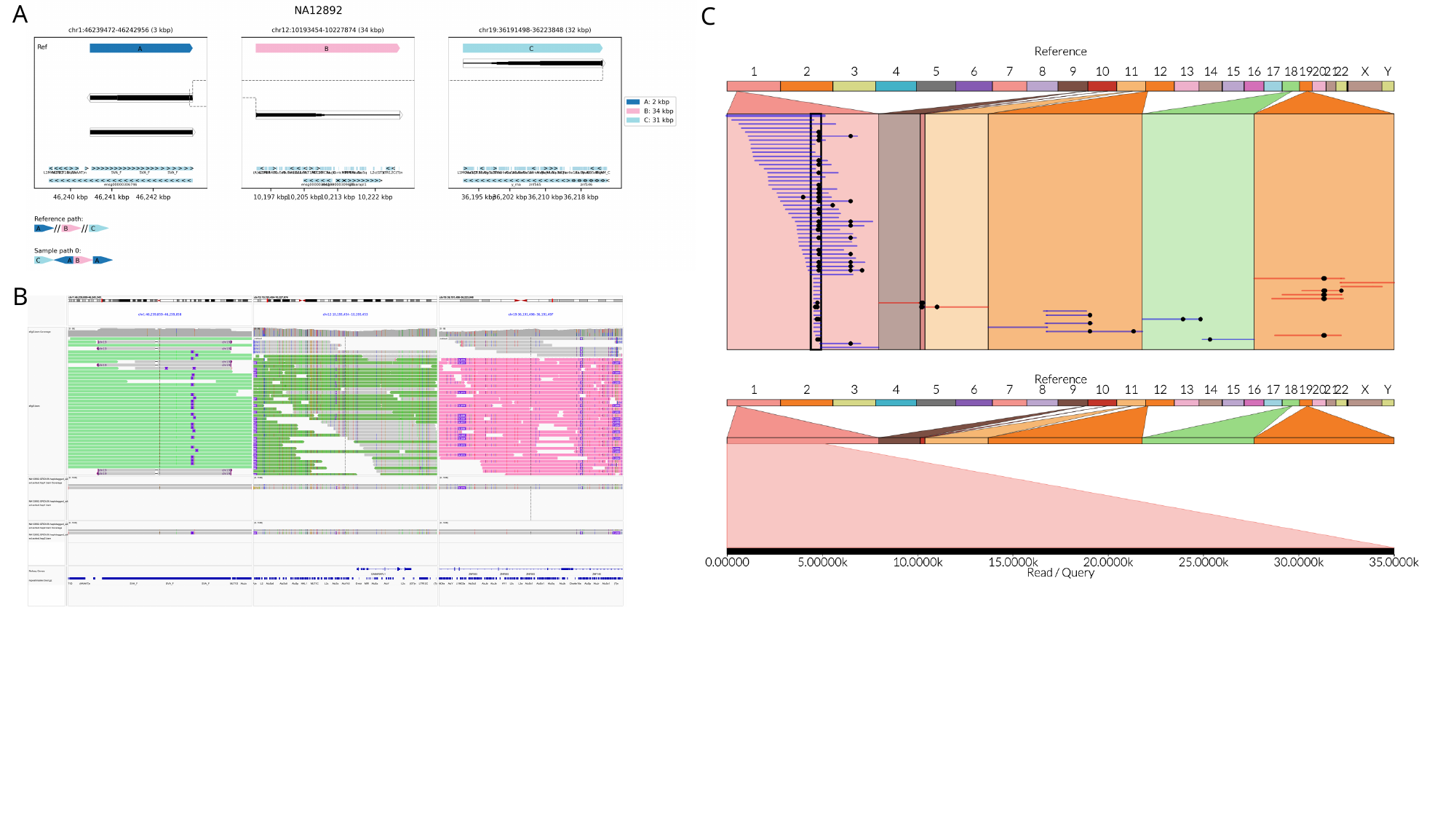

A
C
B

### Slide 13
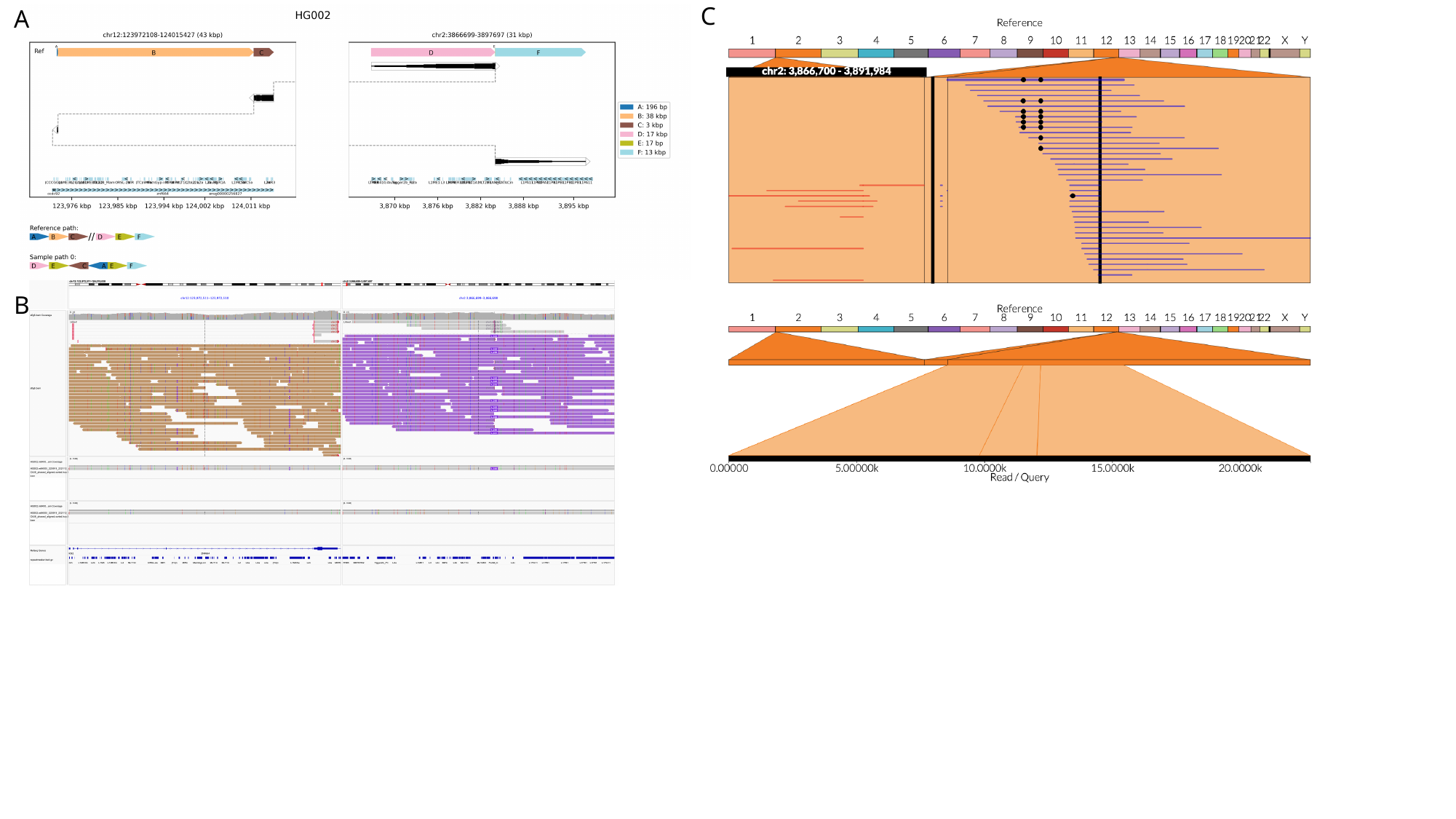

C
A
B

### Slide 14
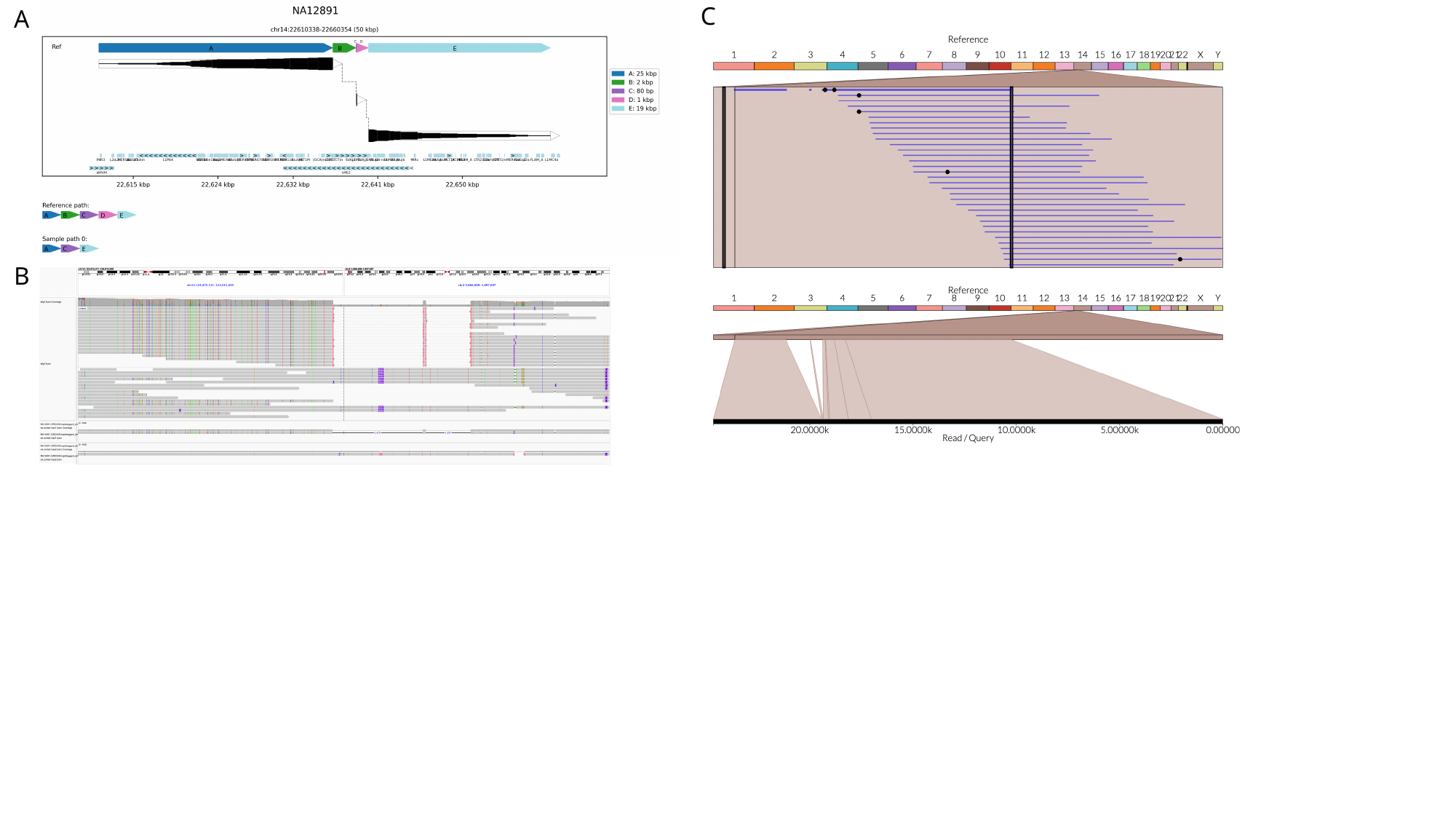

C
A
B

### Slide 15
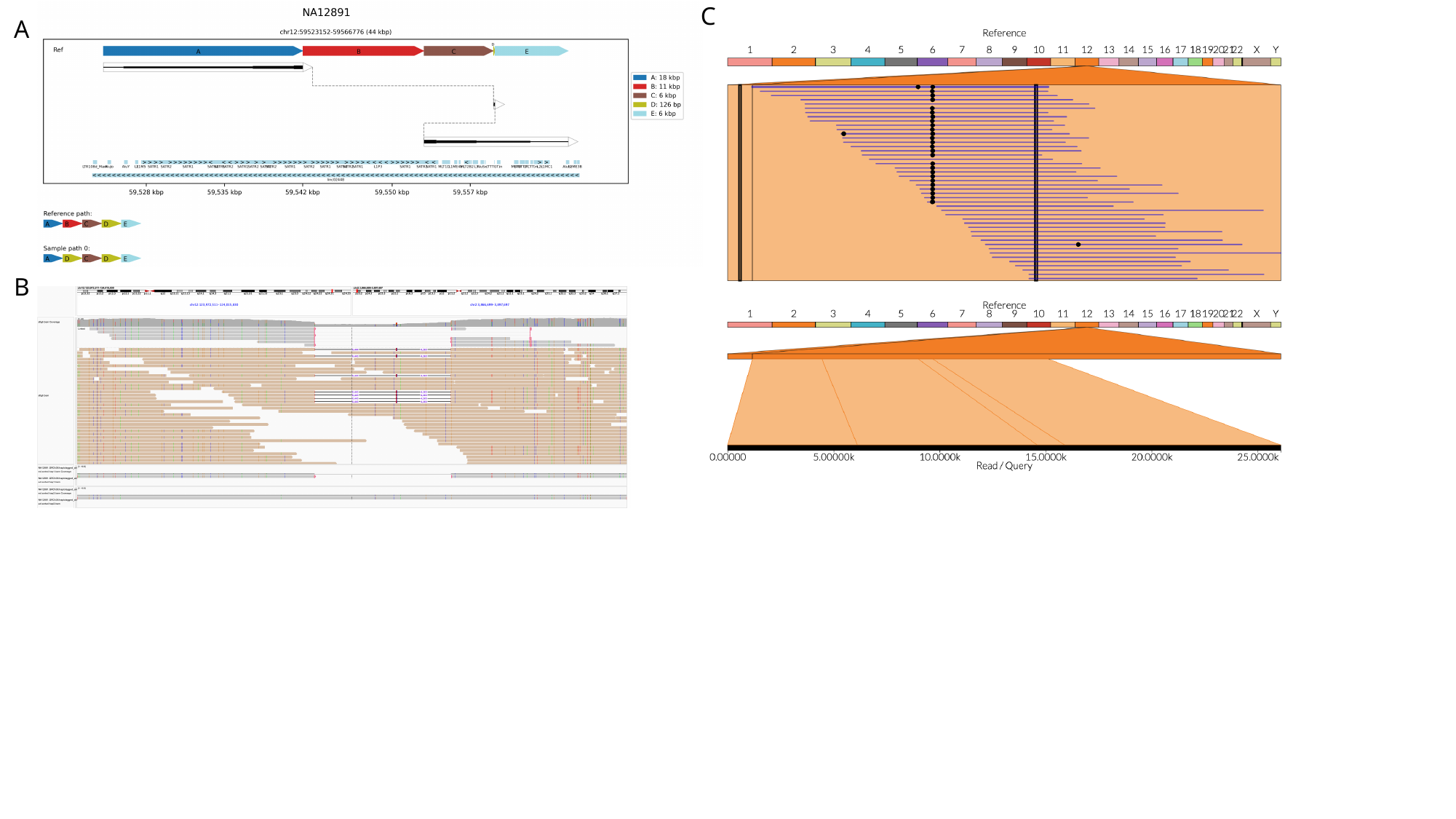

C
A
B

### Slide 16
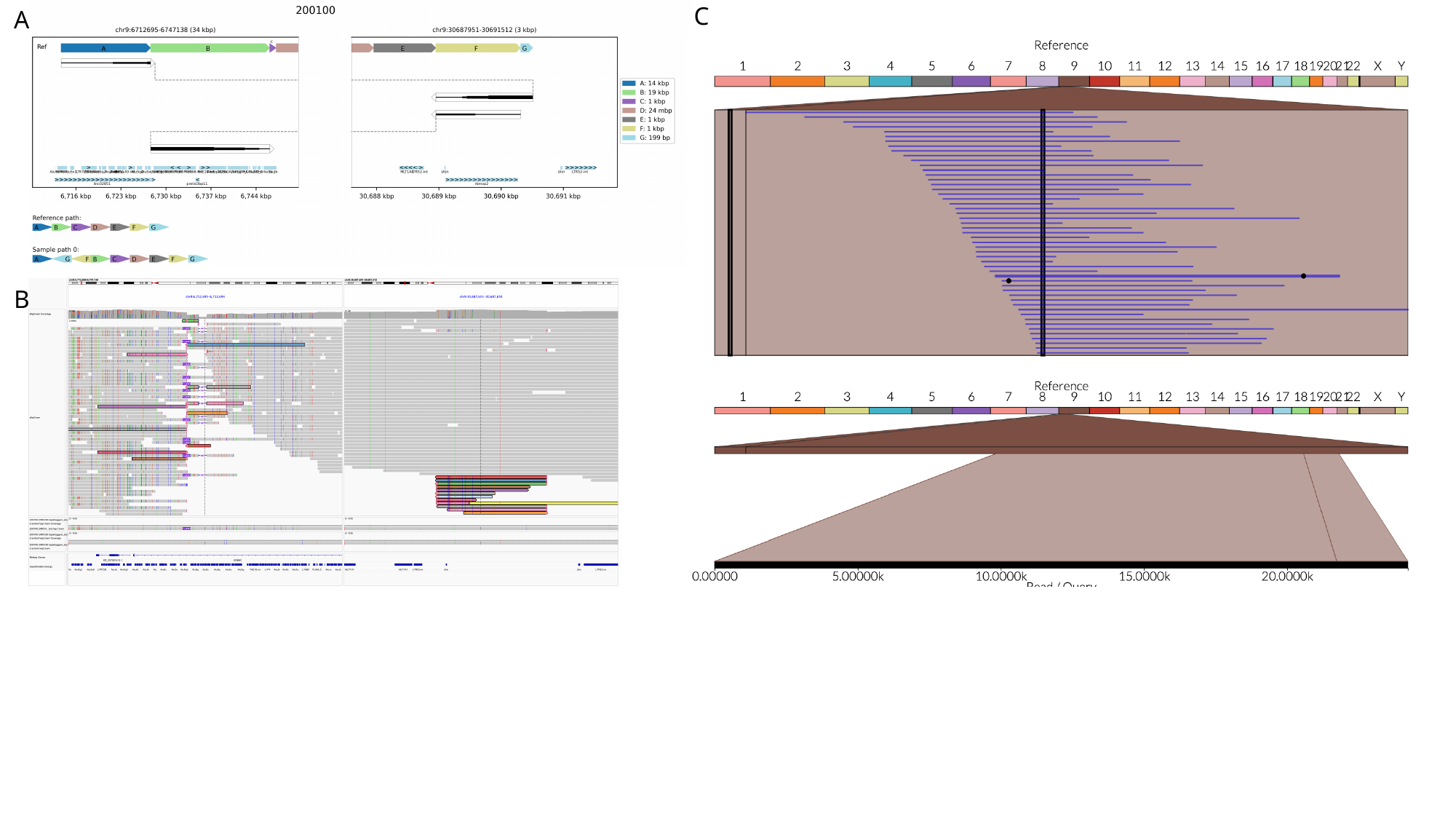

C
A
B

### Slide 17
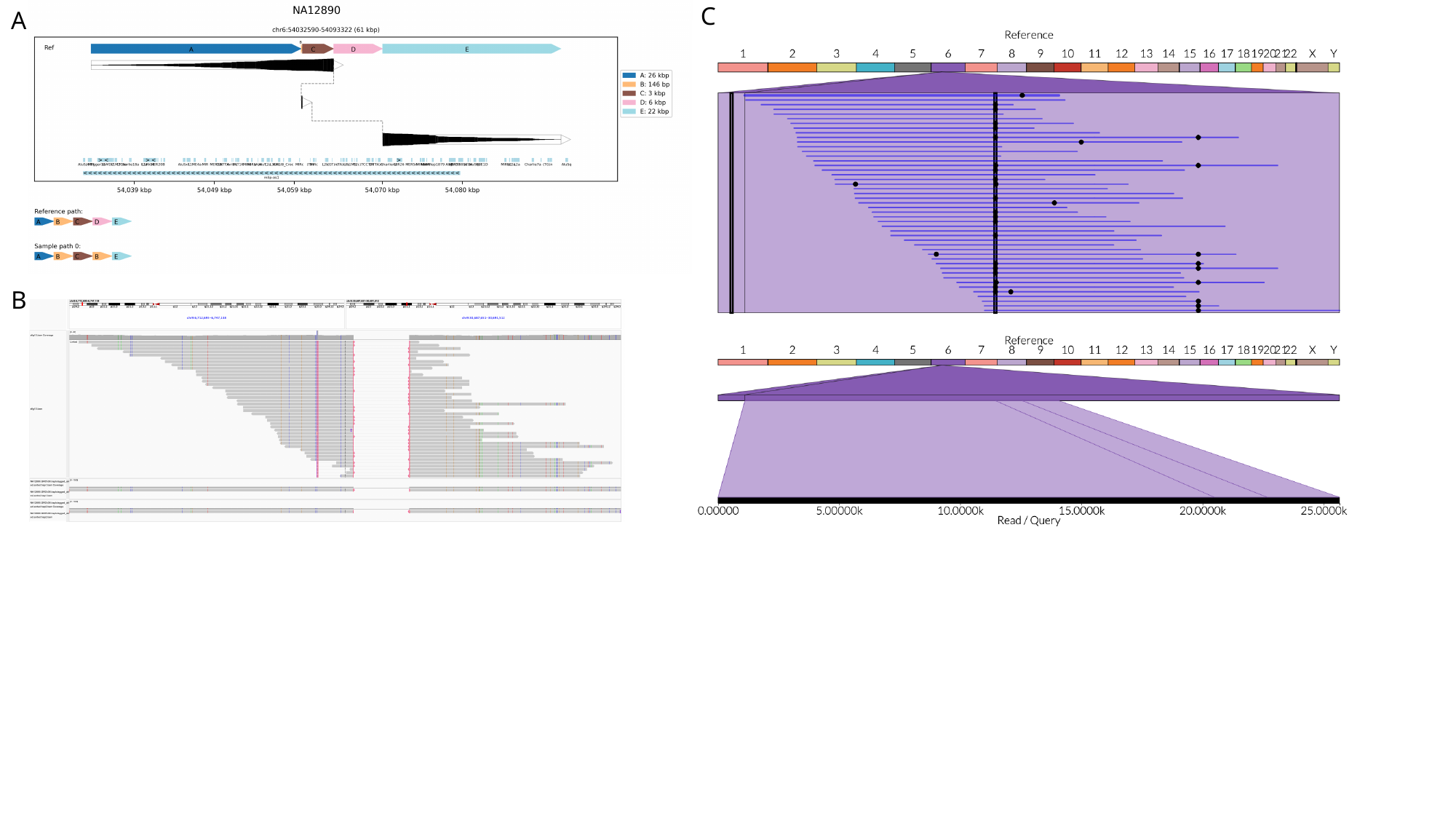

C
A
B

### Slide 18
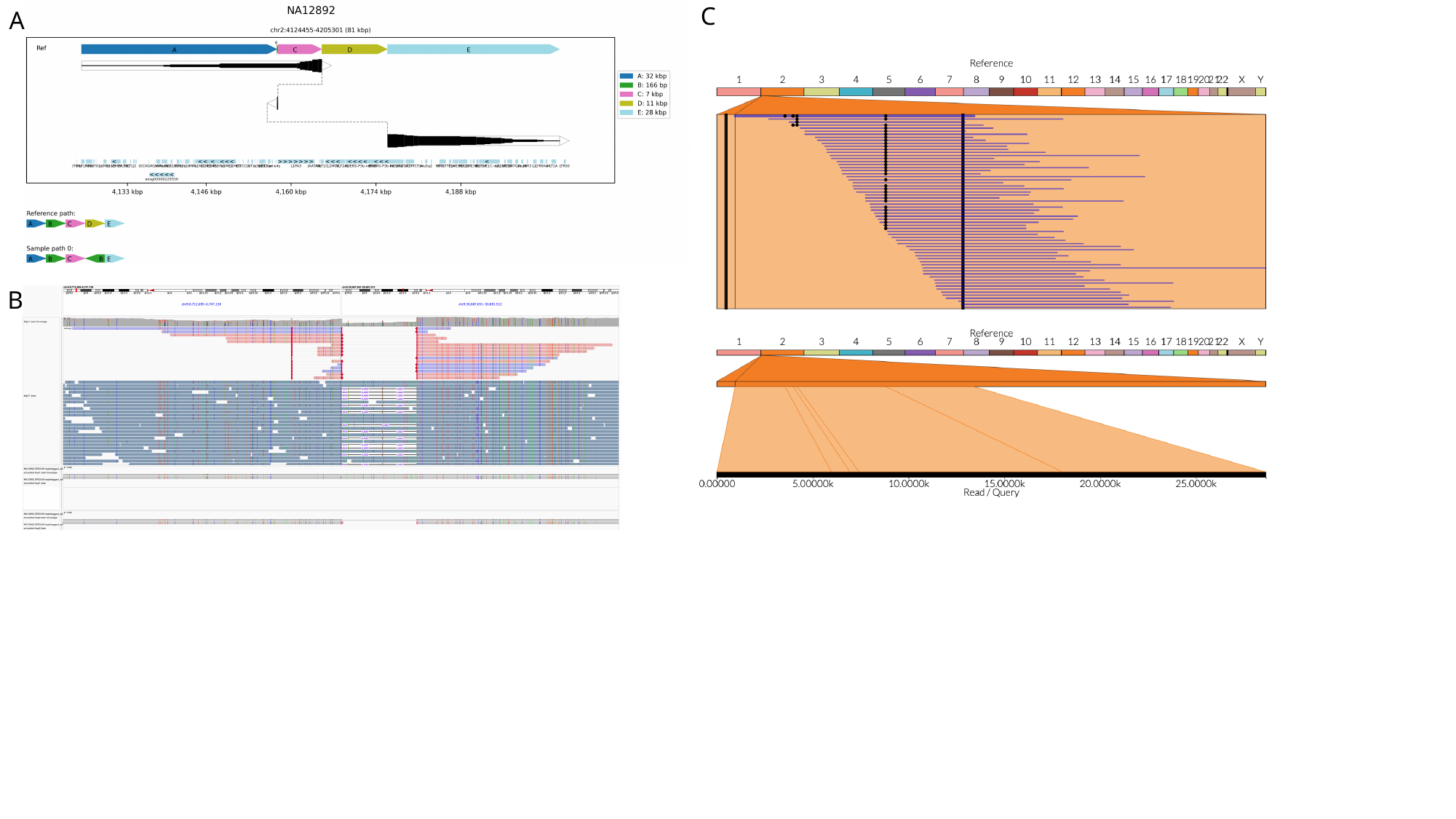

C
A
B

### Slide 19
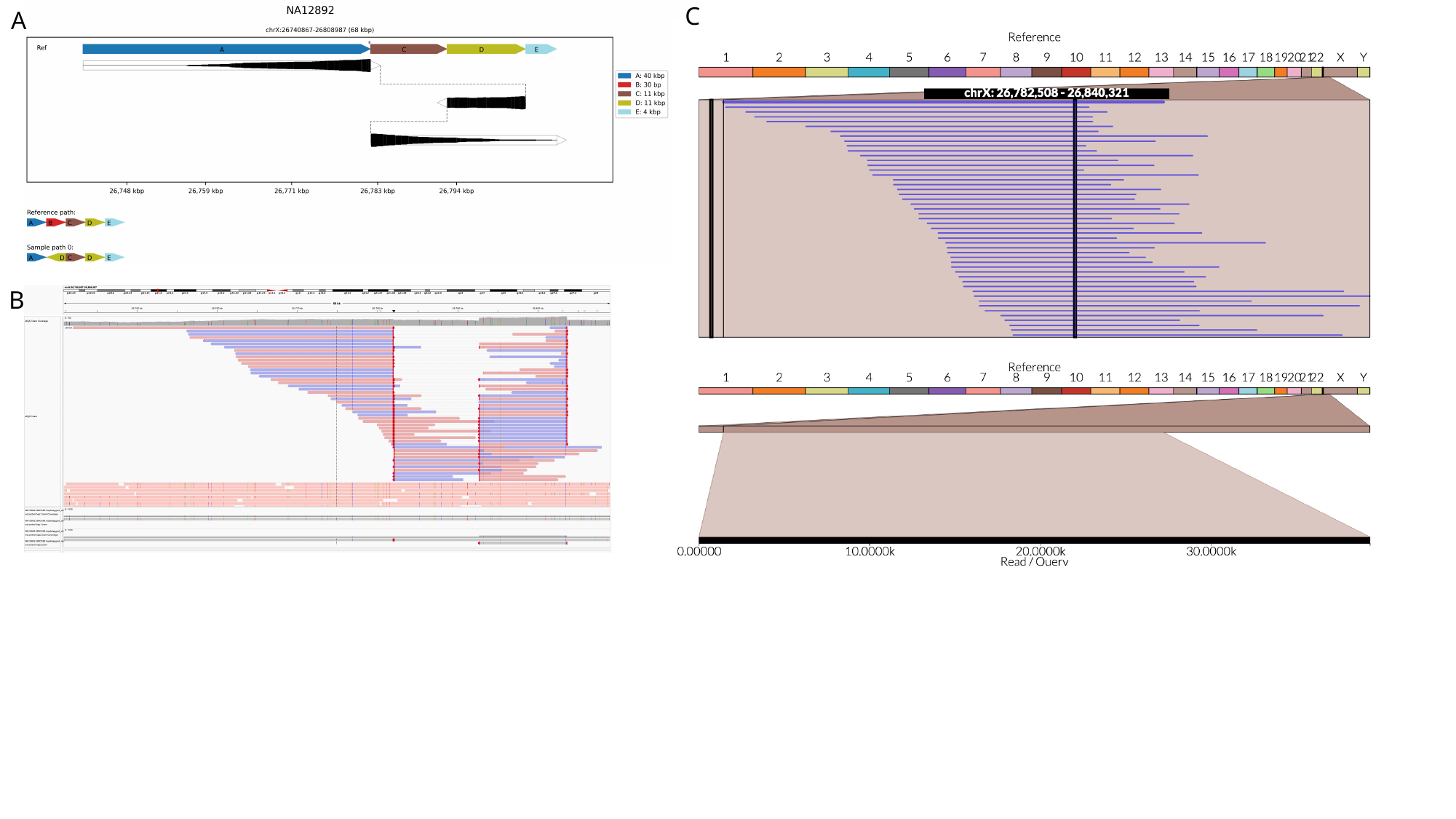

C
A
B
