## Supplementary Note 1 for "Complex structural variant visualization with SVTopo"

### Supplementary Note 1: SVTopo algorithm details.

##### Identifying putative genomic breaks:

SVTopo uses phased high-accuracy long-read alignments to identify breakends in the sample genome, relative to the reference genome. Genomic coordinates where multiple alignments are soft-clipped by the read aligner indicate extensive disagreements between the sample and reference genomes, which often indicate novel adjacencies of genomic sequences in the sample. In SVTopo, only reads with good mapping quality (MAPQ>=20) and at least 100 bases of clipping are considered as evidence for a structural breakend. If at least two alignments share the same clipping location (within a 10 bp margin of aligner error), the coordinate is considered a putative structural variant breakend. The breakend coordinates and supporting alignments are thus associated and stored together.

##### Connecting genomic breaks

Genomic breaks that share at least two supporting alignments are identified as associated in a one-to-one fashion and conceptually interpreted as edges in an undirected graph. Additional edges can be constructed using phasing information if multiple genomic breaks are found in the same phase block and haplotype. Cases where a break is identified with support from alignments that are clipped only on the upstream side, and where a following break has support from alignments only clipped on the downstream side can be paired as two distant ends of the same structural rearrangement if the phase of the supporting alignments matches. These are annotated as derived from phase, as the inferred connections have lower confidence than those with directed alignment evidence.

##### Defining networks of connected breaks

The lists of one-to-one connections of break locations are modeled as a mixed set of distinct graphs. To disentangle these events, a depth-first search algorithm is applied iteratively with randomly selected start locations. Each set of nodes that can be connected to a start location is stored as a new independent event graph and skipped in choosing the next random start. The resulting set of event graphs represents the complex structural rearrangement break locations in the sample genome.

##### Finding event start coordinates

To determine the order of the break locations as they occur in the sample requires first determining the most probably start locations of each rearrangement. If a single break coordinate in the event graph has no connection on the upstream side, it is considered the most likely correct start location. If no break coordinate matches this start definition, a search is performed through the graph to determine if the removal of any phased connections alters a break coordinate to match that definition. If such a phase connection is identified that connection is considered erroneous and is removed to allow use of that break coordinate as the start.

If multiple possible start coordinates are found by either of these methods, the first (in reference space) is selected as the event start. If neither of these methods finds any likely start coordinate, the first of the break coordinates in the complex event graph is selected.

##### VCF support

SVTopo can be supported using output from the sawfish SV caller. If sawfish is run with the --report-supporting-reads option, the resulting JSON file associating variant IDs with VCF entries is used by SVTopo to associate alignments with variant break locations, and to connect break locations together. Each read that is indicated by sawfish to correspond to a break coordinate is scanned and the alignment with clipping that most closely matches the break coordinate is assigned as a supporting alignment for that break coordinate.

##### Ordering complex event graphs

Determining the order of complex events from the unordered complex event graph and start coordinate is complicated by the possibility of coordinates appearing multiple times in the same complex event, creating cycles in the graph. Additionally, in some cases and especially where phase-based connections are used, there is also insufficient information to determine a unique best path through the event graph. A specialized graph traversal approach is used to address these complications, based on breadth-first search.

This modified algorithm is based on a key piece of differentiating evidence between types of graph edge. Each edge is labeled as being ‘spanned’ by alignments or ‘unspanned’. A spanned edge is defined as having alignment support where the clipping on one side of the alignment matches one coordinate node and the clipping on the other side of the alignment matches the other coordinate node. An unspanned edge is defined as having read support where one read alignment is clipped at one coordinate node and the next alignment in the read is clipped at the other coordinate node.

The modified breadth-first search algorithm begins with a queue containing the start coordinate and an index of 0. For each iteration, the current queued coordinate is added to a new ordered event graph data structure at the index given. The neighboring coordinates for the current coordinate are extracted from the unordered event graph and split into two categories: those with a single connection to the current coordinate (either spanned or unspanned) and those with bidirectional connections to the current coordinate (both spanned and unspanned), which represent inverted blocks. Coordinates connected to the current one via bidirectional connections are added to the queue and the current queue is added again as well for each, then unidirectionally connected coordinates are added without the extension back to the current coordinate. All are added to a set of already-processed coordinates to avoid being added again to the queue later.

If the final coordinate added to the ordered event graph is bidirectional and represents a backtrack to an upstream coordinate, a final step is taken to re-add the downstream coordinate and finish the inversion.

##### Annotating the ordered event graph edges with alignment support

To fully represent complex structural variation events, the event graph of break locations is annotated with the evidence supporting edges. The possibilities of revisited coordinates and ambiguous ordering incur some complexity in this process as well as the initial ordering process.

The previously identified start coordinate is processed first and added to a new annotated event graph as a genomic block beginning from the start coordinate of the first supporting alignment and ending at the clipping site. A data structure of genomic coordinates with the number of supporting alignments is added to this block to show the alignment coverage support for the block, and a sample order index of 0 is set to indicate that this is the first block of the rearrangement.

The next sample order index in the un-annotated ordered event graph is processed next. If there are multiple coordinates at that index, they are processed in sequence. For each one, a backward search is performed to find unspanned connections to the previous coordinate(s) in the ordered event graph and any such connections are added to the annotated event graph with an empty alignment coverage data structure. Next, a forward search is performed to find spanned connections to next coordinates in the ordered event graph or to the end of alignment support for the current coordinate. If any such connections are found, they are added to the annotated event graph with their supporting alignment coverage and the downstream coordinate is labeled as already being processed. Finally, a search is performed to find unspanned connections between coordinates tied coordinates at the same sample order index in the un-annotated complex event graph and any found connections are added as blocks. An integer value is used to modify the sample order index from the un-annotated graph to the annotated one, as the addition of spanned or unspanned edges as blocks in the graph alters the index used prior to annotation.

The last block is added similarly to the first, with alignments from the clipped coordinate and extending downstream added as an extension to the last coordinate in the ordered event graph. If the last block in the complex event shares a starting break coordinate with another location in the graph, also check for any unspanned backwards connections to the previous end coordinate and add them first.

##### Annotating the orientations of blocks

The final step in building representations of complex SVs is adding the orientations of blocks already annotated with alignment support. The start block and end blocks are assumed to be in forward (reference) orientation, then each subsequent block (from start and from end) is given an orientation based on the previous block’s orientation. If ambiguous connections exist (such as from phasing), orientation is omitted.

##### Filtering

The annotated graphs are filtered using two criteria: sequencing coverage depth and mapping quality. An iterative pass through the BAM file is used to generate a coverage map. The coverage map uses genomic block coordinates from the annotated event graphs as keys, storing a vector of coverage per-position and a vector of extremely low mapping-quality (defined as MAPQ<5) coverage per-position within the block. After the pass through the BAM to fill the coverage map, annotated event graphs are filtered to remove any with coverage greater than the allowed threshold (default 300x, customizable on the command line), or where the fraction of extremely low-MAPQ coverage is above 5%.

##### Output

The output from SVTopo is a JSON file with a list of annotated complex event graphs. Each graph entry contains a list of block definitions, which consist of the region definition with the start chromosome, start coordinate, end chromosome, and end coordinate; the coverage data structure with genomic coordinates and the supporting alignment coverage at each (empty in the case of unspanned connections), the sample order index, and the orientation of the block.

#### Plotting SVTopo events as images with SVTopoVz

##### Filtering

SVTopoVz begins by applying some additional filtering to events. Event graphs with one event block or fewer are skipped, and those that appear to be simple copy number variants (insertions or deletions) are also skipped. These events are same-chromosome variants where two spanned blocks are connected by a single unspanned block in either forward (deletion) or reverse (duplication) order. They may be optionally included.

##### Selecting plot windows

Genomic windows for the plot are then selected using the event blocks in the event graph. All blocks on different chromosomes are given their own plot window. If individual windows grow larger than the customizable maximum window size, they are subdivided additionally. This allows images to scalably represent SVs that span extremely wide genomic regions.

##### Plotting sample block connections

The main feature in each SVTopoVz plot window is a representation of the spanned and unspanned blocks as they appear in the sample. Blocks are shown in reference genome order from left to right, but on sample order from top to bottom.

##### Plotting chain representation of rearrangements

To assist in understanding the rearrangements of genomic blocks, a representation of chained-together shapes is added for the reference order and the sample order of the blocks. The reference chain plot is shown twice: once at the top of the image with the individual blocks shown in reference-proportional sizes and again at the bottom with all blocks shown at the same size. The sample chain plot representing the order of the blocks as observed in the sample is added at the bottom as well, with uniform block sizes and up two paths through the sample if any block have ambiguous ordering. The number of blocks in the sample chain plot may differ from the reference plot if any blocks are duplicated or deleted. Sizes of the blocks are shown in a legend for the plot.
